# A genetic toolkit of visible mutants for *Caenorhabditis briggsae*

**DOI:** 10.64898/2026.11.19.767928

**Authors:** Robert Johnsen, Allan Mah, Bavithra Thillainathan, Takao Inoue, Michael Ailion, Shahla Gharib, B. Nagagireesh, Dharitri Sarma, Astha Tejpal, David L. Baillie, Paul W. Sternberg, Bhagwati P. Gupta

**Affiliations:** Department of Molecular Biology and Biochemistry, Simon Fraser University, Burnaby, BC V5A 1S6, Canada; Division of Biology and Biological Engineering, California Institute of Technology, Pasadena, CA 91125, USA; Department of Biology, McMaster University, Hamilton, ON L8S 4K1, Canada; Department of Biochemistry, University of Washington, Seattle, WA 98195, USA

**Keywords:** *Caenorhabditis briggsae*, *Caenorhabditis elegans*, forward genetics, genetic linkage map, complementation, genetic markers, levamisole resistance, nicotinic acetylcholine receptor, comparative genetics, synteny

## Abstract

*Caenorhabditis briggsae* is a widely used model for comparative genetic studies involving *C. elegans*. Genetic analysis in *C. briggsae* depends on mutations with clear, easily recognizable phenotypes, yet most mutants reported so far have been mentioned only as names and linkage groups, without underlying experimental data. This study reports 183 *C. briggsae* mutants belonging to 11 phenotypic classes, which represent the most comprehensive collection to date. Almost all mutants were recovered in EMS screens of the reference strain AF16. Our analysis revealed that 121 mutations define 55 loci and 62 remain unassigned. A combination of complementation tests, linkage tests, two- and three-factor crosses, and polymorphism-based mappings helped place 51 loci and 15 further mutations to the six linkage groups. We report the supporting data, map distances, linkage groups, and corrections to several earlier assignments. Twenty-three loci correspond to molecularly identified orthologs of *C. elegans* genes, including six for which we report genetic, rescue, or sequence evidence here. The levamisole-resistant mutants fall into at least three complementation groups, two of which correspond to *Cbr-unc-38* and *Cbr-unc-29* (levamisole-sensitive nicotinic acetylcholine receptor α and non-α subunit, respectively). Transgene rescue experiments identify *Cbr-dpy-1* (mucin-like), *Cbr-sma-6* (TGFβ type receptor kinase), and *Cbr-unc-4* (homeodomain transcription factor), genes that have long been in use as genetic markers. In all cases, molecularly identified genes are located on the same chromosome as their *C. elegans* orthologs, although gene order within chromosomes differs. The strains, map, and underlying data provide a resource for genetic and comparative studies in *Caenorhabditis* nematodes.

## Introduction

*Caenorhabditis briggsae* and *C. elegans* diverged tens of millions of years ago (Stein *et al*. 2003; Cutter 2008), yet the two species exhibit nearly indistinguishable development, behavior and anatomy (Gupta *et al*. 2007). The sequencing and chromosomal assembly of the *C. briggsae* genome revealed that most genes remain on the same chromosome in the two species, although gene order within chromosomes has been extensively rearranged (Stein *et al*. 2003; Hillier *et al*. 2007; Stevens *et al*. 2022; O’connor *et al*. 2026). These features make *C. briggsae* a valuable model for investigating the molecular genetic basis of conserved developmental and physiological processes, as demonstrated by comparative studies of sex determination (Kelleher *et al*. 2008; Guo *et al*. 2013; Chen *et al*. 2014), vulval development (SEETHARAMAN *et al*. 2010; Sharanya *et al*. 2012; Sharanya *et al*. 2015; Chamberlin *et al*. 2020; Jhaveri *et al*. 2025), dauer formation (Inoue *et al*. 2007), and lifespan and stress responses (Amrit *et al*. 2010), all of which depended on *C. briggsae* mutants.

Forward genetic experiments require mutations that cause easily observed phenotypes to facilitate study of gene function, to map new mutations, to follow chromosomes in crosses, and to construct mutant strains. Existing molecular resources for *C. briggsae* include an SNP-based genetic map (Hillier *et al*. 2007), chromosomal-level assembly (Stevens *et al*. 2022; O’connor *et al*. 2026), indel and snip-SNP markers for bulk segregant mapping (Koboldt *et al*. 2010), an SNP microarray (Zhao *et al*. 2010), introgression mapping (Sharanya *et al*. 2015), and genome editing with TALENs and CRISPR/Cas9 (Wei *et al*. 2014; Culp *et al*. 2020). A large collection of mutants with easily observed phenotype, by contrast, has never been described in detail. Earlier, we reported a list of about 50 loci with linkage groups and easily observed mutant phenotypes, but no supporting data were provided (Gupta *et al*. 2007). Several of these loci have since been used as mapping markers by our laboratories and others (Inoue *et al*. 2007; Kelleher *et al*. 2008; Guo *et al*. 2009; Koboldt *et al*. 2010; Sharanya *et al*. 2012; Yan *et al*. 2012; Chen *et al*. 2014; Sharanya *et al*. 2015; Le *et al*. 2017), and some carry the names of *C. elegans* orthologs (for example *Cbr-dpy-1*, *Cbr-sma-6*, *Cbr-unc-4* and *Cbr-unc-22*) for which the supporting data have not been described.

Here we present the collection of *C. briggsae* mutations in full detail. The set comprises mutants isolated in EMS screens using the reference strain AF16, along with a few alleles contributed by other laboratories. We define loci by complementation, molecular cloning, and map position, assign them to linkage groups, and present the linkage tests along with two and three-factor crosses that position the genes. Some of the earlier map assignments have been corrected in light of molecular mapping and other linkage data. We report genetic, rescue, and allele sequencing results for several *C. elegans* orthologs, including two subunits of the levamisole-sensitive nicotinic acetylcholine receptor, describe the complementation groups, and drug responses of levamisole-resistant mutants. We also report initial characterization of several egg-laying defective mutants not reported elsewhere. Some of the loci described here have been molecularly characterized in a parallel study (Lyu *et al*. 2026). Finally, we compare the chromosomal locations of identified genes in *C. briggsae* and *C. elegans*. Together, these data serve as a substantial resource for *C. briggsae* genetics.

## Materials and Methods

### Strains and culture

*C. briggsae* strains were maintained at 20 °C on NGM plates seeded with *Escherichia coli* OP50 (Brenner 1974; Wood 1988). The wild strain AF16 was used to generate mutants and as the wild-type control for other assays. The wild strain HK104 was used for polymorphism-based mapping. All loci, alleles and strains, along with mutant phenotypes, crosses, rescue and molecular data for each locus, are listed in Supplementary File S1. Allele numbers include every mutation recovered and assigned to a locus. For loci with many alleles, only representative alleles were kept as frozen stocks; Supplementary File S1 lists the strains that are available. Eight alleles were contributed by other researchers: 2 *ge* alleles (A. Fodor and D. Riddle), 3 *gu* alleles (H. M. Chamberlin), 2 *nm* alleles (E. S. Haag), and *mf104* (M.-A. Félix).

### Mutagenesis and screens

AF16 hermaphrodites were mutagenized with 25 mM ethyl methanesulfonate (EMS, Sigma Aldrich) for 3 - 4 h following the *C. elegans* protocol (Brenner 1974). Worms were washed with M9 buffer and placed singly on plates. In some of the screens, we used 1% agarose in place of agar to minimize burrowing. F2 progeny were screened for easily observed phenotypes. Candidates were cloned and confirmed in subsequent generations.

For levamisole-resistant mutants, 1 mM levamisole hydrochloride (Sigma Aldrich) containing agar plates were used to screen F2 progeny of mutagenized worms. Animals alive and moving after 3 - 4 min were picked singly, and their descendants were re-tested two to three generations later. Eight of 12 candidates bred true and one of the false positives was a twitcher. Two other *lev* alleles were isolated independently.

Mutants were outcrossed to AF16 two to five times before confirming phenotypes and further characterization, and all lines were established from single animals. At least 250,000 haploid genome sets were screened in three laboratories (PWS, DLB, and BPG). Screens for egg-laying defective (Egl) and vulva mutants have been described (Sharanya *et al*. 2012; Sharanya *et al*. 2015), as have screens for dauer-formation mutants (Inoue *et al*. 2007). Egl (Egg laying defective, bloated with retained eggs) and Pvl (protruding vulva) phenotypes were scored on plates (100 or more per strain). Vulval morphology of Lin class and male tail of Muv animals were examined under Nomarski (15-20 animals per genotype).

### Genetic nomenclature

Gene and allele names follow *C. elegans* conventions (Tuli *et al*. 2018). Loci without a molecular identity are named by phenotypic class and reference allele, as are mutations not assigned to loci. Molecularly identified genes carry the Cbr prefix. Several informal class names were used in earlier work, including Cby (chubby) for Dpy-like, Mip (movement impaired) for Unc-like, Rot (rotator) for Rol-like and Sml for Sma-like loci (Gupta *et al*. 2007; Inoue *et al*. 2007). We retire these names to avoid confusion with *C. elegans* nomenclature and with other *C. briggsae* loci. For example, *cby-3* has been used for both *dpy(sy5039)* and an unrelated X-linked allele, *bd101* (CGC strain PB101, isolated by S. E. Baird), now identified as an allele of *Cbr-dpy-6* (Lyu *et al*. 2026). Supplementary File S1 lists old and new names.

### Complementation tests and definition of loci

Complementation tests were carried out by crossing hermaphrodites homozygous for one mutation with males heterozygous for the other and scoring cross progeny. In some cases, a GFP marker such as *myo-2::GFP* or *myo-3::GFP* was used to distinguish between self and cross progeny. For X-linked mutations, F1 cross-progeny hermaphrodites were scored, because F1 males carry only the maternal X chromosome.

A mutation was assigned to a locus if it (i) failed to complement another mutation; (ii) carried a mutation in, or was rescued by, an identified gene; (iii) complemented other alleles of its phenotypic class on the same linkage group, or of every locus of its class if unmapped; or (iv) was the only mutation of its class on its linkage group. Previously described loci follow the cited papers. Loci defined by (iv) have not been tested for complementation and could be allelic to unmapped mutations of the same class or to genes with a different mutant phenotype. Mutations that met none of these criteria are listed as not assigned to loci.

### Linkage tests, two and three-factor mappings

Linkage to a chromosome was tested against marker mutations of known linkage group. LGI was marked by *Cbr-lin-11*, *Cbr-unc-38(sy5440)*, or *Cbr-pry-1(sy5353)*, LGII by *Cbr-unc-4* or *Cbr-sma-6*, LGIII by *Cbr-dpy-1* alleles, LGIV by *Cbr-unc-22 or Cbr-dpy-4*, LGV by *Cbr-unc-68(sa997)*, and the X chromosome by the phenotype of hemizygous F1 males. In clonal assays, F2 progeny of trans heterozygotes (*a* +/+ *b*) showing one phenotype but not the other were picked singly, and their progeny scored for the second phenotype. For genes that are unlinked, two-thirds of such clones are expected to segregate the second phenotype. When F2 animals showing one phenotype are scored directly for the second, we can expect one-quarter to show it without linkage. For X-chromosome linkage, mutant hermaphrodites were crossed with wild-type males and cross-progeny males were examined for the mutant phenotype.

Map distances were calculated from F2 counts using one of the following six equations, chosen according to the cross design. p is a recombination frequency, reported in map units (mu; 1 mu = 1% recombination), except in Equation 6, where it is already in mu. Equation 1 is described earlier (Brenner 1974). Equations 2 and 3 follow from Brenner’s point that recombination can be estimated from one recombinant class, and equation 4 inverts his expression for the fraction of recombinant heterozygotes among parental animals picked from a trans heterozygote. Equations 5 and 6 are the standard estimates for the designs indicated.

**(1)** p = 1 - √(1 - 2R), for F2 progeny of hermaphrodites heterozygous for two mutations in cis (a b/+ +), when all four phenotypic classes can be scored. R is the fraction of recombinant progeny. **(2)** p = 1 - √(1 - 4f), when double mutants cannot be distinguished from one of the single-mutant classes (e.g., when a paralyzed Unc phenotype masks Dpy), only the other single mutant class is used. f is the fraction of all F2 progeny in that class. **(3)** p = 1 - √(1 - 3f), when Dpy F2 animals are not counted in a cross involving *dpy* and *unc* mutations, because the Dpy and Dpy Unc classes cannot be clearly distinguished. f is the fraction of non-Dpy F2 animals that are Unc. **(4)** p = f / (2 - f), for heterozygotes carrying the two mutations in trans (a +/+ b), F2 A non-B animals are picked singly, and f is the fraction whose progeny segregated B. As mentioned above, this equation follows Brenner’s explanation. Animals with parental phenotype (say, A) can be either a +/a + or recombinant heterozygotes a +/a b. In this case, the fraction of the recombinant class is 2p(1-p) / ((1-p)^2^ + 2p(1-p)), which simplifies to 2p / (1+p) and gives p = f / (2-f). **(5)** p = r / t, when recombinant chromosomes can be counted directly, F2 animals homozygous for one marker are tested for the second marker. r is the number of recombinant chromosomes and t the number of chromosomes tested (Table 3). **(6)** p = f x d, provides the distance of the mutation from one marker, where f is the fraction of crossovers that occurred on that side of the mutation and d is the distance (mu) between the markers. Used in three-factor crosses, in which heterozygous hermaphrodites carrying two markers in cis and a mutation of interest in trans (a + b/+ m +) were allowed to self. The fraction of each recombinant class carrying the mutation places it relative to the markers.

### Polymorphism-based mapping

Mutants were crossed to HK104 males, and DNA from pools of F2 mutant and wild-type animals was genotyped for indel and snip-SNP polymorphisms (Koboldt *et al*. 2010) or hybridized to an SNP microarray (Zhao *et al*. 2010). For *unc(sy5422)*, single F2 animals were genotyped for SNPs on the AF16 x HK104 recombinant inbred map (Hillier *et al*. 2007) by R. D. Miller, Washington University St. Louis (personal communication). Indels used in this study are described (Koboldt *et al*. 2010) (Supplementary File S1).

### Transgenic rescue

*C. elegans* or *C. briggsae* DNA together with a *myo-2*::GFP marker was microinjected into the gonads of *C. briggsae* animals, and stable extrachromosomal lines were established from GFP-expressing F1s. *C. elegans* fosmids WRM0632cF10 (*unc-38*) and WRM0625cF12 (*unc-29*) were injected into *lev(sy5440)*, *lev(sy5437)* and *lev(sy5436)* (Table 6; Supplementary File S1). The *C. elegans* cosmid M01E10 (10 ng/µl), which carries *dpy-1* (M01E10.2), was injected into *dpy(ge12)*, and two lines were obtained. A 6 kb *C. briggsae* genomic fragment spanning *Cbr-sma-6* open reading frame was amplified and cloned in pCR-XL-TOPO. The fragment was injected into *sma(sy5148)*. The *C. elegans unc-4* genomic plasmid pNC4-21 (provided by D. M. Miller) was injected into *unc(sy5341)*, and three stable lines were obtained. Strain and array names of all transgenic lines are listed in Supplementary File S1.

For levamisole assays, transgenic animals carrying arrays were identified by GFP fluorescence of microinjection marker, transferred with non-transgenic siblings to plates containing 1 mM levamisole and scored for paralysis. Rescue was scored when GFP-expressing animals were paralyzed and their siblings lacking GFP were not.

### Drug sensitivity

Stock solutions of levamisole hydrochloride (1 M in water), aldicarb (1 M in 70% ethanol), and ivermectin (10 mg/ml in DMSO) were added to NGM before pouring. Alternatively, a 10x drug stock was spread on NGM plates and allowed to diffuse into the agar overnight before use.

For Lev mutants, adult animals were placed on 1 mM levamisole and paralysis of the head and body was scored under a stereomicroscope. Growth of Lev mutants and AF16 control was compared on plates containing levamisole (1 or 2 mM) or aldicarb (1 or 5 mM), and was scored qualitatively (>100 worms per plates per strain; 3 or more independent batches). Growth of *C. briggsae* AF16 and *unc(sy5422)*, and *C. elegans unc-119* animals was compared on plates containing 0, 5, 10, or 20 ng/ml ivermectin and scored qualitatively (20-30 animals per assay per strain; two to three independent batches).

### Genome resources and sequence analysis

Physical positions of molecularly identified *C. briggsae* genes refer to the CGC2 assembly of an isogenic derivative of AF16 (O’connor *et al*. 2026). Positions were taken from the cb4 Wormbase gene models that were lifted to CGC2 with Liftoff (Shumate and Salzberg 2021; O’connor *et al*. 2026) (https://github.com/AndersenLab/CGC2_genome_MS). As an independent check, we aligned the first and last 150 bp of each gene in the cb4 assembly (Ensembl Metazoa release 116) to the CGC2 genome sequence (CaeNDR release 20261102; https://caendr.org) with minimap2 (Ross *et al*. 2011; Li 2018; Yates *et al*. 2022). Positions in the cb4 assembly are given in Table 5 for comparison. Physical positions (WBcel235 assembly), genetic map positions, alleles, and phenotypes of *C. elegans* genes are from WormBase release WS298, queried through WormMine (Sternberg *et al*. 2024). Chromosome lengths were taken from the same assemblies. Strain records and allele designations are listed in the CGC database (https://cgc.umn.edu). Positions are given as the lower gene coordinate. Positions of mutations estimated by SNP-microarray mapping (Zhao *et al*. 2010) refer to the earlier cb3 assembly and mark the peak of the linkage signal, which can lie up to about 2 Mb from the gene (Zhao *et al*. 2010). Since then, the cb4 assembly moved part of the cb3 sequence to other chromosomes and ordered or reoriented large parts of the rest (Ross *et al*. 2011). Therefore, cb3 positions cannot be compared directly with cb4 or CGC2 positions.

Protein sequences were obtained from UniProt (Uniprot 2025): *C. briggsae* UNC-38 (A8XXX4), *C. elegans* UNC-38 (Q23022), *Drosophila melanogaster* nicotinic acetylcholine receptor α1 (P09478), and the muscle acetylcholine receptor α1 subunits (CHRNA1) of zebrafish (Q98880), mouse (P04756) and human (P02708).

Full-length sequences were aligned with Clustal Omega 1.2.4 (Sievers *et al*. 2011). MAFFT 7.505 (Katoh and Standley 2013) gave an identical alignment of the Cys-loop region, which required no gaps. Vertebrate residues are also given in mature-protein numbering, lacking the 20-residue signal peptide.

### Molecular Biology

Sequences of all primers are given in Supplementary Table S1. Coding exons were amplified from mutant genomic DNA and sequenced. *Cbr-unc-38* was amplified with primers that cover all exons. All 11 exons of *Cbr-unc-29* were amplified with primers in two independent rounds from PCR experiments. Protein positions refer to the 510-residue CBG20491.1 model of Cbr-UNC-38 (UniProt A8XXX4).

For quantitative RT-PCR, total RNA was extracted from mixed-stage AF16 and *lev(sy5437)* animals and reverse transcribed. *Cbr-unc-29* transcripts were amplified and normalized to *Cbr-rsp-5*. Results are means ± SD of three independent biological replicates.

### Data analysis, figures, and tables

Map distances and exact binomial confidence (exact binomial 95% confidence intervals, CI) were calculated for the observed fraction (Clopper and Pearson 1934), and both limits were converted to map distances with the same expression, as done previously in *C. elegans* (Sigurdson *et al*. 1986). Calculations use the counts in Table 3 and File S1 and were done in Python 3.13 with SciPy 1.18 library (Virtanen *et al*. 2020b; Virtanen *et al*. 2020a) using a script written by Claude Opus 5.5 (Anthropic Inc.) (Supplementary File S2). Distances are in map units (mu; 1 mu = 1% recombination). Figures 2 and 4 were plotted from data in Tables 1 - 6, Supplementary File S1, and known *C. elegans* gene positions (Wormbase release WS298) using Claude Opus 5.5. Other figures were generated in Microsoft PowerPoint and Adobe Illustrator. Figure panels were exported as PNG and TIFF files and assembled into final figures in PowerPoint.

### Data availability

Some of the strains are available from the *Caenorhabditis* Genetics Center (CGC), and the others are available on request. Supplementary File S1 contains the complete list of mutations, strains, phenotypes, and other relevant details.

## Results

### *C. briggsae* mutations isolated from forward genetic screens of AF16

The collection of strains comprises 183 mutations in 11 phenotypic classes (Table 1, Figure 1B, D, Supplementary File S1). 121 mutations define 55 loci (Figure 1D), including 23 with confirmed molecular identities. The remaining 62 mutations are not assigned to loci, mostly because they have not been tested against other mutations of the same class. Figure 1C shows, for each linkage group, the molecularly identified genes, other loci and unassigned mutations. The largest phenotypic classes are Dumpy (Dpy; 72 mutations, 61 belonging to 13 loci, and 11 unassigned mutations), Uncoordinated (Unc; 29 mutations, 6 loci and 23 unassigned), Lineage-defective (Lin; 30 mutations, 27 belonging to 16 loci, 3 unassigned) and Egg-laying defective (Egl; 18 mutations, 4 belonging to 3 loci, 14 unassigned).

**Figure 1.**
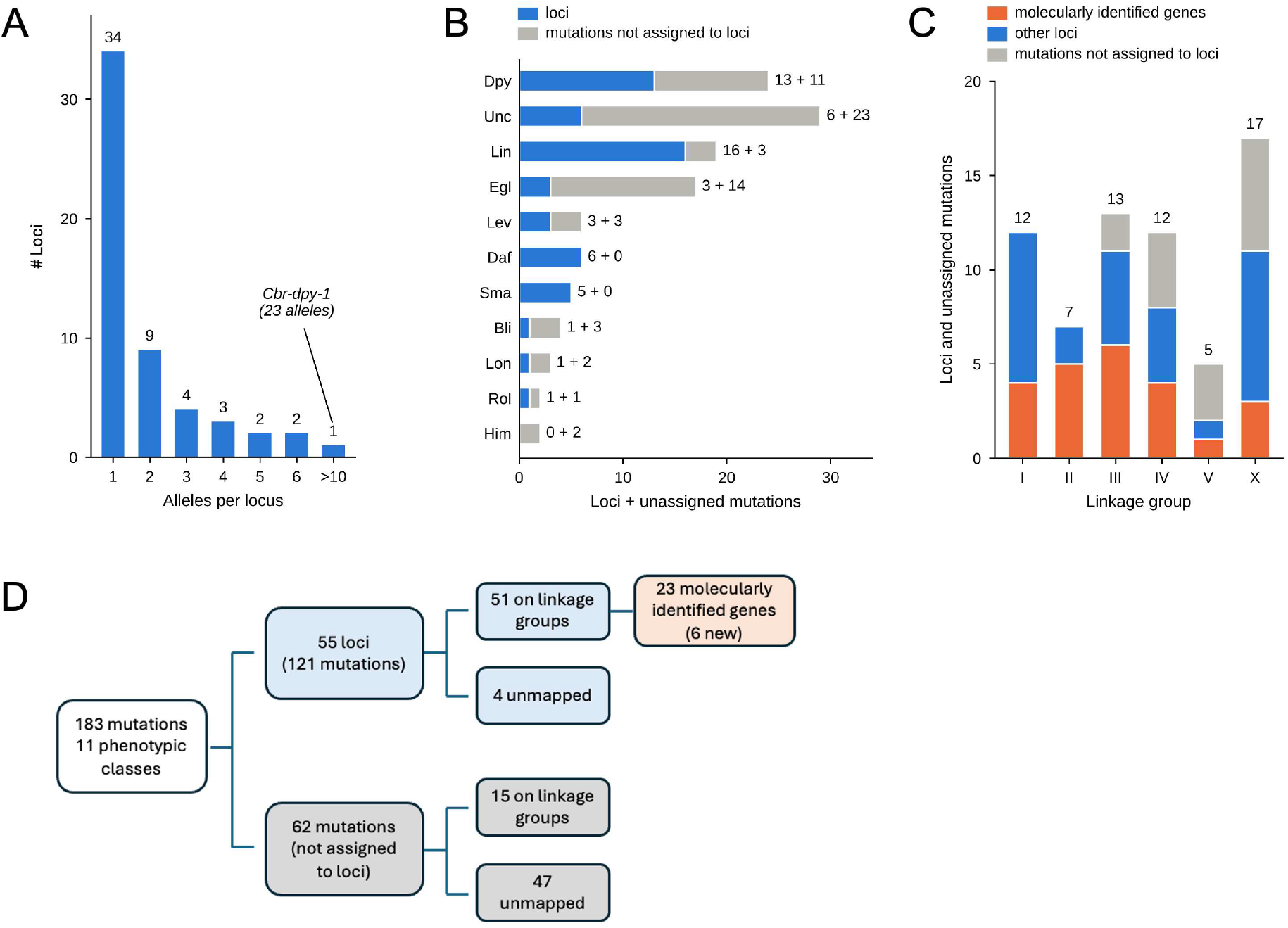
Overview of the *C. briggsae* mutant collection. (A) Histogram showing number of alleles per locus. Mutations not assigned to loci are excluded. Most loci are defined by a single allele; *Cbr-dpy-1* has 23. (B) Number of loci (blue) and of mutations not assigned to loci (grey) in each phenotypic class. (C) Stacked bars show loci and unassigned mutations on each linkage group: molecularly identified genes (orange), other loci (blue) and mutations not assigned to loci (grey). (D) Classification of the 183 mutations. Loci are defined by molecular identity, complementation, published analyses or map position (see Materials and Methods), and mutations that meet none of these criteria are not assigned to loci.

**Table 1.**
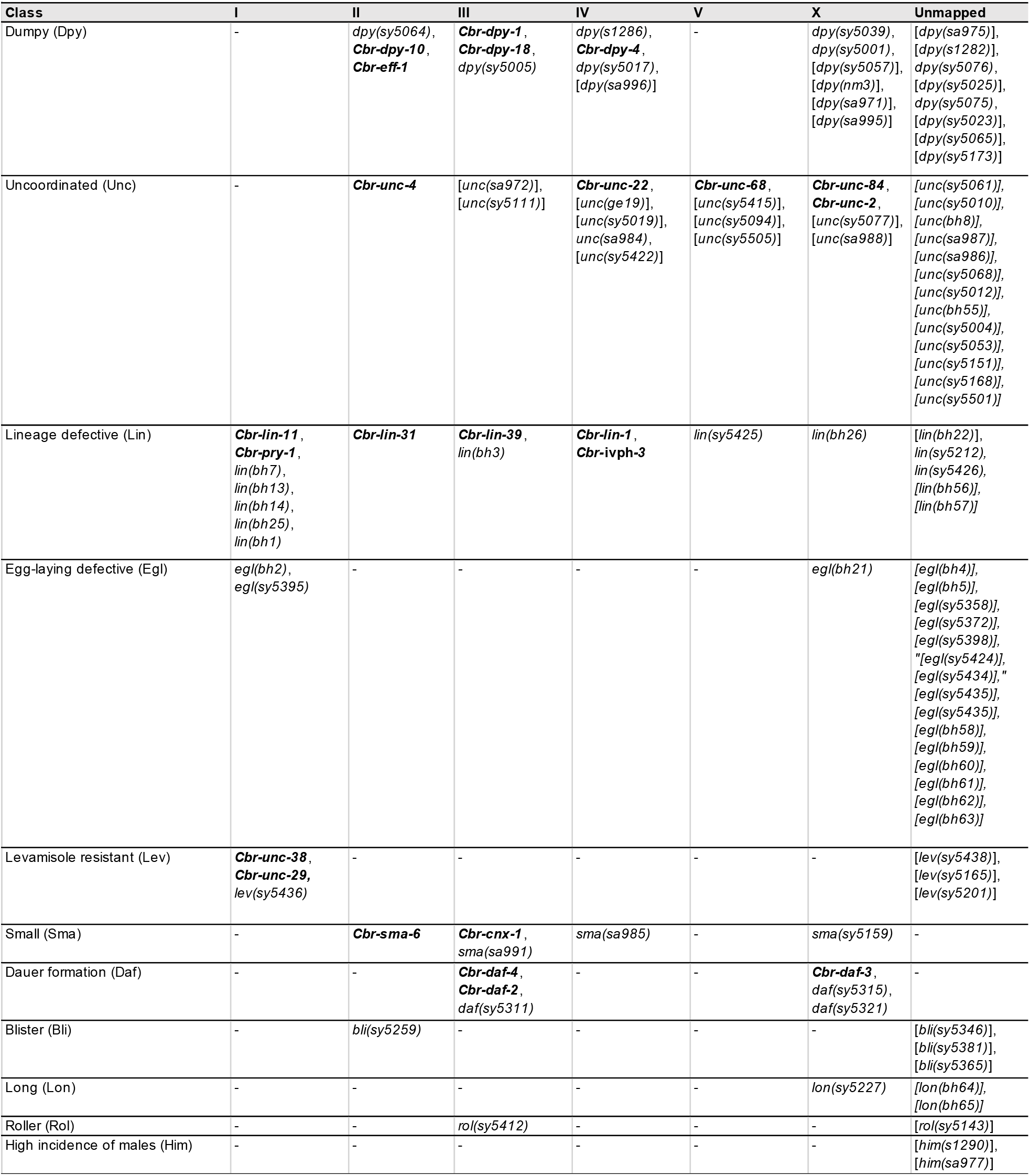
*C. briggsae* mutations and loci of different phenotypic classes placed on linkage group. Molecularly identified genes are in bold. Mutations not assigned to loci are in square brackets. See Supplementary File S1 for details.

| Class | I | II | III | IV | V | X | Unmapped |
|---|---|---|---|---|---|---|---|
| Dumpy (Dpy) | - | *dpy(sy5064)*, ***Cbr-dpy-10***, ***Cbr-eff-1*** | ***Cbr-dpy-1***, ***Cbr-dpy-18***, *dpy(sy5005)* | *dpy(s1286)*, ***Cbr-dpy-4***, *dpy(sy5017)*, [*dpy(sa996)*] | - | *dpy(sy5039)*, *dpy(sy5001)*, [*dpy(sy5057)*], [*dpy(nm3)*], [*dpy(sa971)*], [*dpy(sa995)*] | [*dpy(sa975)*], [*dpy(s1282)*], *dpy(sy5076)*, [*dpy(sy5025)*], *dpy(sy5075)*, [*dpy(sy5023)*], [*dpy(sy5065)*], [*dpy(sy5173)*] |
| Uncoordinated (Unc) | - | ***Cbr-unc-4*** | [*unc(sa972)*], [*unc(sy5111)*] | ***Cbr-unc-22***, [*unc(ge19)*], [*unc(sy5019)*], *unc(sa984)*, [*unc(sy5422)*] | ***Cbr-unc-68***, [*unc(sy5415)*], [*unc(sy5094)*], [*unc(sy5505)*] | ***Cbr-unc-84***, ***Cbr-unc-2***, [*unc(sy5077)*], [*unc(sa988)*] | *[unc(sy5061)],* *[unc(sy5010)],* *[unc(bh8)],* *[unc(sa987)],* *[unc(sa986)],* *[unc(sy5068)],* *[unc(sy5012)],* *[unc(bh55)],* *[unc(sy5004)],* *[unc(sy5053)],* *[unc(sy5151)],* *[unc(sy5168)],* *[unc(sy5501)]* |
| Lineage defective (Lin) | ***Cbr-lin-11***, ***Cbr-pry-1***, *lin(bh7)*, *lin(bh13)*, *lin(bh14)*, *lin(bh25)*, *lin(bh1)* | ***Cbr-lin-31*** | ***Cbr-lin-39***, *lin(bh3)* | ***Cbr-lin-1***, ***Cbr-*****ivph-3** | *lin(sy5425)* | *lin(bh26)* | [*lin(bh22)*], *lin(sy5212),* *lin(sy5426),* *[lin(bh56)],* *[lin(bh57)]* |
| Egg-laying defective (Egl) | *egl(bh2)*, *egl(sy5395)* | - | - | - | - | *egl(bh21)* | *[egl(bh4)],* *[egl(bh5)],* *[egl(sy5358)],* *[egl(sy5372)],* *[egl(sy5398)],* *"[egl(sy5424)],* *[egl(sy5434)],"* *[egl(sy5435)],* *[egl(sy5435)],* *[egl(bh58)],* *[egl(bh59)],* *[egl(bh60)],* *[egl(bh61)],* *[egl(bh62)],* *[egl(bh63)]* |
| Levamisole resistant (Lev) | ***Cbr-unc-38***, ***Cbr-unc-29,*** *lev(sy5436)* | - | - | - | - | - | [*lev(sy5438)*], [*lev(sy5165)*], [*lev(sy5201)*] |
| Small (Sma) | - | ***Cbr-sma-6*** | ***Cbr-cnx-1***, *sma(sa991)* | *sma(sa985)* | - | *sma(sy5159)* | - |
| Dauer formation (Daf) | - | - | ***Cbr-daf-4***, ***Cbr-daf-2***, *daf(sy5311)* | - | - | ***Cbr-daf-3***, *daf(sy5315)*, *daf(sy5321)* | - |
| Blister (Bli) | - | *bli(sy5259)* | - | - | - | - | [*bli(sy5346)*], [*bli(sy5381)*], [*bli(sy5365)*] |
| Long (Lon) | - | - | - | - | - | *lon(sy5227)* | *[lon(bh64)],* *[lon(bh65)]* |
| Roller (Rol) | - | - | *rol(sy5412)* | - | - | - | [*rol(sy5143)*] |
| High incidence of males (Him) | - | - | - | - | - | - | [*him(s1290)*], [*him(sa977)*] |

Other classes include Levamisole-resistant (Lev; 10 mutations, 7 belonging to 3 loci, 3 unassigned), Dauer-formation defective (Daf; 6 loci), Small (Sma; 7 mutations belonging to 5 loci), Blister (Bli; 1 locus and 3 unassigned mutations), Long (Lon; 1 locus and 2 unassigned), Roller (Rol; 1 locus and 1 unassigned) and High incidence of males (Him; 2 unassigned mutations). See Supplementary Table S2 for information about the publication status of mutations.

Thirty-four of the 55 loci are represented by a single allele, whereas *Cbr-dpy-1* has 23 alleles and *dpy(sy5076)* and *dpy(sy5017)* have six each (Figure 1A). Because many of the loci are defined by single alleles, and many mutations remain unassigned, the screens were not saturated, a conclusion that was also noted in one of our earlier studies (Sharanya *et al*. 2012). The expectation is that many more genes should be recoverable in future genetic screens with easily observable mutant phenotypes.

### Phenotypic classes

#### Dpy

The Dpy class includes *Cbr-dpy-1*, whose animals are short with a thickened posterior. These include strains such as *Cbr-eff-1(s1281)*, *dpy(sy5039)*, *dpy(sy5025)*, and *dpy(sy5075)* that exhibit a strong Dpy phenotype. Several alleles also exhibit a rolling movement, which is typical of cuticle collagen mutants in *C. elegans*. *dpy(sy5001)* is a medium Dpy that rolls to the right, *dpy(sa975)* is a weak Dpy with a dominant roller phenotype, *dpy(sy5023)* is a strong Dpy that rolls weakly to the left, and some *dpy(sy5005)* animals roll during backward movement. *Cbr-dpy-1* alleles were recovered more often than any other locus, suggesting that the gene is a large mutational target, as in *C. elegans*.

#### Unc

The 29 Unc mutations exhibit a wide range of movement defects. Animals carrying *Cbr-unc-22* alleles show a twitching phenotype, similar to *C. elegans unc-22* mutants (Moerman and Baillie 1979). *unc(sy5341)* animals move forward normally but cannot move backward, as do *C. elegans unc-4* mutants. Other Unc mutants include *unc(ge19)*, *unc(sy5019)* and *unc(sa984)* that are coilers and *unc(sa988)*, *unc(sy5094)* and *unc(sy5077)* that exhibit kinker phenotype. Two other Unc strains, *unc(sy5422)* and *unc(sy5415),* give rise to animals that are small and nearly paralyzed. Although *unc(sy5415)* form dauers, *unc(sy5422)* animals do not.

#### Lin and Egl

The Lin and Egl classes comprise mutants recovered in screens for defects in egg laying and vulval morphology. The mapped Lin loci and three of the Egl loci have been described (Seetharaman *et al*. 2010; Sharanya *et al*. 2012; Sharanya *et al*. 2015; Chamberlin *et al*. 2020; Jhaveri *et al*. 2023). They are included here for completeness along with new Egl mutants.

The Lin class includes vulvaless (Vul) and multivulva (Muv) mutants that exhibit defects in vulval precursor cell (VPC) induction and cell lineage (Seetharaman *et al*. 2010; Sharanya *et al*. 2012; Sharanya *et al*. 2015; Jhaveri and Gupta 2023). Two of the strains, *lin(bh56)* and *lin(bh57)*, were not described earlier. The animals show abnormal vulval invagination and adults are Pvl. Besides vulval defects, some of the Lin mutants exhibit male tail phenotypes. *Cbr-pry-1(sy5353)* males have ectopic anterior rays and fused rays and no hook, and *Cbr-ivph-3(sy5392)* males lack rays and have a defective hook and crumpled spicules. *Cbr-lin-1(sa993)* males show normal rays and hooks, but some have crumpled spicules, as do some *Cbr-lin-31(sy5344)* males. These defects agree with the mating defects of Muv males noted earlier (Sharanya *et al*. 2015).

The phenotypic analyses of Egl mutants including penetrance, vulval anatomy and responses to serotonin and fluoxetine of *egl(bh2)*, *egl(sy5395)* and *egl(bh21)* have been described (Sharanya *et al*. 2012). The remaining 14 Egl strains show variable egg-laying defects and are mostly not fully penetrant. Six are also Pvl and four are also Unc (Supplementary File S1). Detailed analysis of developing vulva and other components of egg-laying system was not performed.

#### Remaining phenotypic classes

The 10 Lev mutations cause an Unc phenotype and resistance to levamisole. Their complementation groups, drug responses, and molecular identification are described below. The Sma class includes *Cbr-sma-6*, whose animals resemble strong Dpy mutants, and four other loci, *Cbr-cnx-1*, *sma(sa991)*, *sma(sy5159)*, and *sma(sa985)*. *bli(sy5259)* animals show blisters over the entire body with complete penetrance. *lon(sy5227)* animals are long and thin. *rol(sy5412)* animals are right-handed rollers at all stages.

The six Daf loci were characterized including the molecular identities of *Cbr-daf-2*, *Cbr-daf-3*, and *Cbr-daf-4* (Inoue *et al*. 2007). We include them here because they are part of genetic screens and serve as markers on LGIII and X.

### Assignment of loci to linkage groups

Of the 55 loci, 51 are assigned to linkage groups: 12 to LGI, 7 to LGII, 11 to LGIII, 8 to LGIV, 2 to LGV, and 11 to the X chromosome (Table 1, Figure 1C, D). A further 15 mutations not assigned to loci are also mapped: 2 on LGIII, 4 on LGIV, 3 on LGV, and 6 on the X chromosome. Every linkage group carries at least one molecularly identified gene, and all except LGV carry at least three (see below), which tie it to a *C. briggsae* chromosome. LGV is defined by *Cbr-unc-68(sa997), unc(sy5094),* and *unc(sy5415)*, all of which were mapped by indels (Table 2). *Cbr-unc-68(sa997)*, a ryanodine receptor homolog (Lyu *et al*. 2026), had earlier been placed on chromosome V by polymorphism-based mapping (Koboldt *et al*. 2010), as had the four other LGV mutations, *lin(sy5425)*, *unc(sy5094)*, *unc(sy5415)*, and *unc(sy5505)*, with SNP or indel markers (Koboldt *et al*. 2010; Zhao *et al*. 2010; Sharanya *et al*. 2012).

**Table 2.**
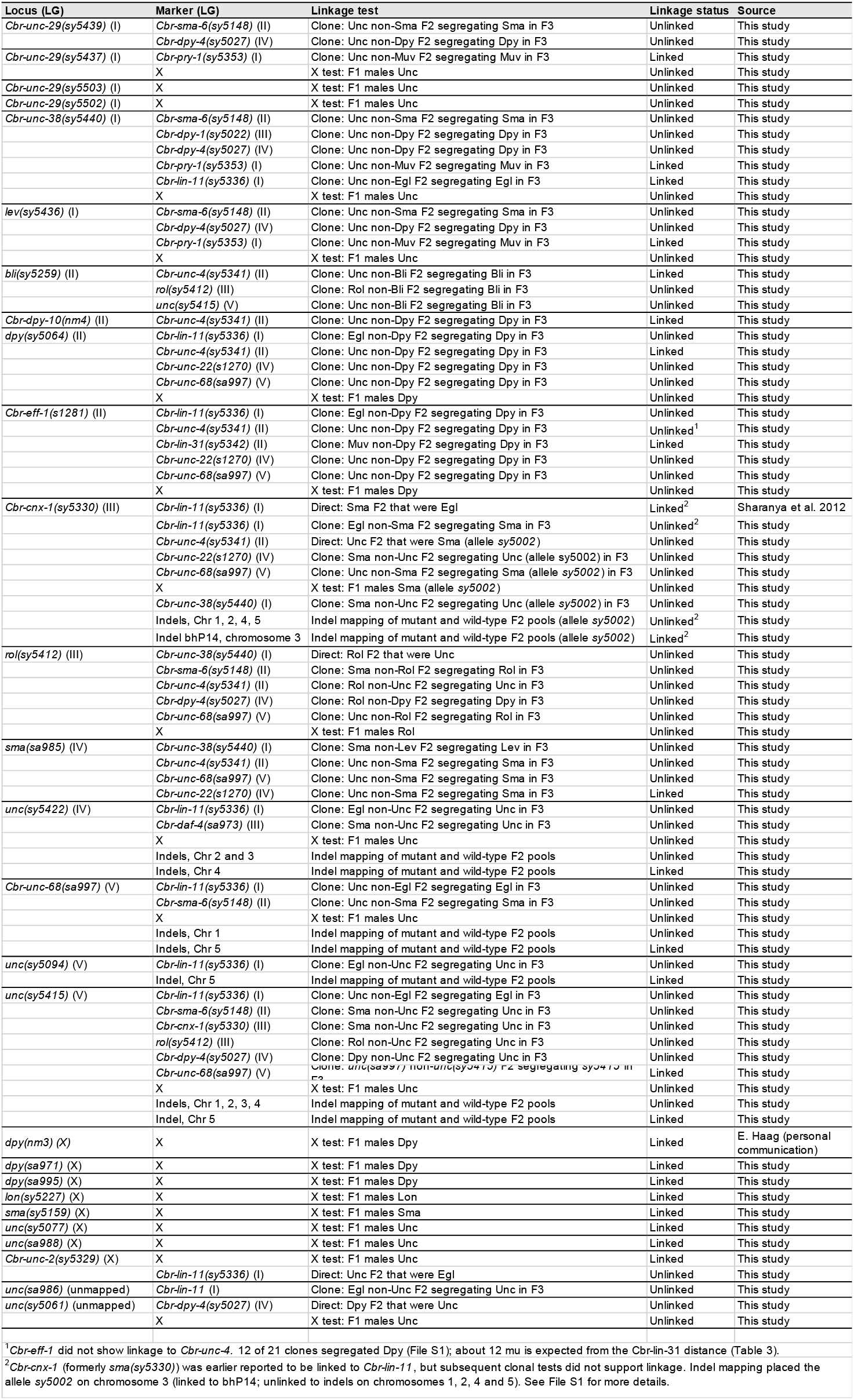
Linkage tests used to assign loci to linkage groups or to exclude linkage groups. Loci are grouped by linkage group. See Materials and Methods and Supplementary File S1 for details. Chr, chromosome.

Most other assignments are based on linkage to marker mutations. Table 2 shows linkage tests, including those that exclude other linkage groups, and Table 3 the map distances. About 30 loci and mutations have also been mapped using polymorphic markers (Koboldt *et al*. 2010; Zhao *et al*. 2010; Sharanya *et al*. 2012; Sharanya *et al*. 2015; Jhaveri and Gupta 2023). Additional details are below.

**Table 3.** Map distances for loci. Counts are F2 phenotypic classes (wild-type / the two single-mutant classes / double mutant) for cis heterozygotes unless stated. A dash means that the cross shows linkage but gives no reliable distance. See Materials and Methods for more details.

| LG | Interval | Counts | n | Eq. | mu | 95% CI | Notes |
|---|---|---|---|---|---|---|---|
| I | *Cbr-unc-38(sy5440) - Cbr-pry-1(sy5353)* | 768 / 98 / 53 / 205 | 1124 | 1 | 14.5 | 12.2 - 17.0 | A clone test gave 18.8 mu (9.6 - 32.1; 12/38 Unc non-Muv F2 clones segregated Muv; File S1) |
| I | *Cbr-unc-38(sy5440) - Cbr-lin-11(sy5336)* | 0/42 Unc non-Egl F2 clones segregated Egl in F3 | 42 | 4 | 0 | 0.0 - 4.4 | See also the three-factor cross (Table 4) |
| I | *Cbr-unc-29(sy5437) - Cbr-pry-1(sy5353)* | 7/22 Unc non-Muv F2 clones segregated Muv in F3 | 22 | 4 | 18.9 | 7.4 - 37.8 |  |
| I | *lev(sy5436) - Cbr-pry-1(sy5353)* | 12/34 Unc non-Muv F2 clones segregated Muv in F3 | 34 | 4 | 21.4 | 11.0 - 36.5 | Side not determined |
| II | *Cbr-sma-6(sy5148) - Cbr-unc-4(sy5341)* | 437 / 46 / 55 / 109 | 647 | 1 | 17.1 | 13.9 - 20.8 |  |
| II | *Cbr-dpy-10(nm4) - Cbr-unc-4* | 324 / 33 / 35 / 91 | 483 | 1 | 15.2 | 11.8 - 19.4 |  |
| II | *dpy(sy5064) - Cbr-unc-4* | 500 WT, 66 Unc non-Dpy | 566 | 3 | 19.4 | 14.8 - 25.0 | Dpy animals not counted. A clone test gave 5.0 mu (0.6 - 17.9; File S1). |
| II | *bli(sy5259) - Cbr-unc-4* | 14/53 Unc non-Bli F2 clones segregated Bli in F3 | 53 | 4 | 15.2 | 8.3 - 25.3 |  |
| II | *Cbr-lin-31(sy5342) - Cbr-unc-4* | 20 of 50 recombinants from *Cbr-sma-6 Cbr-unc-4 / Cbr-lin-31(sy5342)* hets | 50 | 6 | 6.8 | 4.5 - 9.4 | Share of the 17.1-mu interval (Table 4); 10.3 mu from *Cbr-sma-6* (Sharanya et al. 2015) |
| II | *Cbr-eff-1(s1281) - Cbr-lin-31(sy5342)* | 7/72 Muv non-Dpy F2 clones segregated Dpy in F3 | 72 | 4 | 5.1 | 2.0 - 10.5 | *Cbr-eff-1* side on the chromosome based on its physical position (3.50 Mb) |
| III | *Cbr-dpy-1(s1272) - unc(sa972)* | 871 / 42 / 46 / 276 | 1235 | 1 | 7.4 | 5.9 - 9.1 |  |
| III | *rol(sy5412) - Cbr-cnx-1* | 2/126 F2 clones segregated the second mutation in F3: 1/44 Rol non-Sma (sy5330); 1/49 Rol non-Sma and 0/33 Sma non-Rol (sy5002) | 126 | 4 | 0.8 | 0.1 - 2.9 | Three clone tests with two *Cbr-cnx-1* alleles pooled (File S1) |
| III | *Cbr-cnx-1(sy5002) - unc(sy5111)* | 546 / 20 / 23 / 191 | 780 | 1 | 5.7 | 4.1 - 7.6 | Cross used the *Cbr-cnx-1* allele *sy5002* |
| III | *sma(sa991) - unc(sy5111)* | 564 / 10 / 12 / 20 | 606 | - | - | - | Linked<sup>1</sup> |
| III | *unc(sy5111) - dpy(sy5005)* | 443 / 85 / 87 / 124 | 739 | 1 | 26.9 | 22.9 - 31.4 |  |
| IV | *Cbr-unc-22(s1270) - Cbr-dpy-4(sy5027)* | 547 / 20 / 24 / 196 | 787 | 1 | 5.8 | 4.2 - 7.7 |  |
| IV | *Cbr-unc-22(s1270) - dpy(s1286)* | 485 / 34 / 45 / 135 | 699 | 1 | 12.0 | 9.5 - 15.0 |  |
| IV | *Cbr-unc-22(s1270) - dpy(sy5017)* | 442 / 58 / 137 / 53 | 690 | 2 | 18.5 | 13.8 - 24.5 | Unc non-Dpy class: 58 |
| IV | *Cbr-unc-22(s1270) - Cbr-lin-1(sa993)* | 431 / 83 / 88 / 68 | 670 | 1 | 30.0 | 25.5 - 35.2 |  |
| IV | *Cbr-unc-22(s1270) - sma(sa985)* | 15/40 Unc non-Sma F2 clones segregated Sma in F3 | 40 | 4 | 23.1 | 12.8 - 37.2 | Side not determined |
| IV | *unc(ge19) - Cbr-dpy-4(sy5027)* | 289 / 13 / 13 / 86 | 401 | 1 | 6.7 | 4.4 - 9.8 |  |
| IV | *unc(ge19) - dpy(s1286)* | 659 / 70 / 58 / 173 | 960 | 1 | 14.4 | 12.0 - 17.1 |  |
| IV | *unc(sa984) - Cbr-dpy-4(sy5027)* | 759 / 45 / 24 / 207 | 1035 | 1 | 6.9 | 5.4 - 8.7 |  |
| IV | *unc(sy5019) - Cbr-dpy-4(sy5027)* | 486 / 21 / 120 / 29 | 656 | 2 | 6.6 | 4.1 - 10.2 | Unc non-Dpy class: 21 |
| IV | *unc(sy5019) - dpy(sy5017)* | 267 / 54 / 51 / 91 | 463 | 1 | 26.1 | 21.2 - 31.8 |  |
| IV | *unc(sa984) - dpy(sy5017)* | 402 / 71 / 87 / 137 | 697 | 1 | 26.1 | 22.0 - 30.7 |  |
| IV | *dpy(sa996) - Cbr-unc-22(s1270)* | 577 / 146 / 138 / 100 | 961 | 1 | 36.1 | 31.7 - 40.9 | Position not determined<sup>2</sup> |
| IV | *dpy(sa996) - unc(ge19)* | 810 / 158 / 146 / 137 | 1251 | 1 | 28.3 | 25.1 - 31.8 | Position not determined<sup>2</sup> |
| IV | *dpy(sa996) - unc(sa984)* | 730 / 116 / 99 / 77 | 1022 | 1 | 23.9 | 20.7 - 27.4 | Position not determined<sup>2</sup> |
| IV | *dpy(sa996) - unc(sy5019)* | 433 / 88 / 67 / 53 | 641 | 1 | 28.1 | 23.7 - 33.2 | Position not determined<sup>2</sup> |
| V | *unc(sy5415) - Cbr-unc-68(sa997)* | 15/35 Cbr-unc-68 non-unc(sy5415) clones segregated unc(sy5415) in F3 | 35 | 4 | 27.3 | 15.2 - 43.5 | Side not determined |
| X | *unc(sa988) - dpy(sy5039)* | 2 recombinants / 40 chromosomes | 40 | 5 | 5.0 | 0.6 - 16.9 | From the three-factor cross (Table 4) |
| X | *unc(sa988) - dpy(sy5001)* | 6 recombinants / 76 chromosomes | 76 | 5 | 7.9 | 3.0 - 16.4 |  |
| X | *dpy(sy5039) - dpy(sy5001)* | 1 recombinant / 40 chromosomes | 40 | 5 | 2.5 | 0.1 - 13.2 | From the three-factor cross (Table 4) |
| X | *unc(sy5077) - dpy(sy5001)* | 490 / 46 / 149 / 38 | 723 | 2 | 13.7 | 9.9 - 18.5 | Unc non-Dpy class: 46 |

<sup>1</sup>The double-mutant class is far below expectation, so no distance is given
<sup>2</sup>dpy(sa996) is linked to all four markers, but its position is not determined. The four distances can be reconciled with a single position only if unc(ge19), unc(sa984) and unc(sy5019) lie on the side of Cbr-dpy-4 opposite Cbr-unc-22, which has not been tested.

*unc(sy5422)* is placed on chromosome IV by three independent methods: SNP genotyping of single F2 animals (R. Miller, personal communication), SNP-microarray mapping (Zhao *et al*. 2010), and indel mapping (Koboldt *et al*. 2010) (Table 2). The mutation was earlier assigned to LGIII based on *Cbr-dpy-1(sy5022)* linkage (Gupta *et al*. 2007), which was not supported by subsequent mapping (Table 2). The Dpy mutation *mf104*, a Mos1 insertion in *Cbr-dpy-18* (CBG13195) generated in the Félix laboratory, lies on chromosome III and serves as an LGIII marker (Chen *et al*. 2014; Le *et al*. 2017). The Egl mutation *egl(sy5395)* maps to the left arm of chromosome I by bulk segregant analysis (Sharanya *et al*. 2012). *unc(sa986)* was earlier assigned to LGV based on preliminary results (Gupta *et al*. 2007), but the data could not be independently verified and the mutation is now listed as unmapped. Another *dpy* mutation, *dpy(sa996)*, is placed on LGIV, as previously reported (Gupta *et al*. 2007).

However, linkage tests involving four different mutations produced estimates that disagree with one another (24-36 mu range) (Table 3), precluding us from precise placement of its location.

*Cbr-sma-6*, defined by two alleles, *sy5148* and *sa983* (Supplementary File S1), maps to LGII by linkage to *Cbr-unc-4* (Table 3). Another Sma locus, *Cbr-cnx-1* (formerly *sma(sy5330)*), was used as an LGI marker (Inoue *et al*. 2007; Sharanya *et al*. 2012; Sharanya *et al*. 2015), but four lines of evidence place it on LGIII. First, it is tightly linked to *rol(sy5412)* (0.8 mu; Table 3), which maps to chromosome III (Zhao *et al*. 2010). Second, indel mapping of *sy5002* (another allele of *sy5330*) showed linkage to chromosome III and none to chromosomes I, II, IV and V (Table 2). Third, we observed no linkage to the LGI markers *Cbr-lin-11* and *Cbr-unc-38*, or to markers on LGII, LGV and the X chromosome (Table 2). And, fourth, *sy5330* is a missense mutation in *Cbr-cnx-1*, which lies at 9.37 Mb on chromosome III (Lyu *et al*. 2026). The earlier LGI assignment depended on a single cross with *Cbr-lin-11* (Sharanya *et al*. 2012), but a subsequent test showed no linkage (18 of 26 Egl clones segregated Sma; Table 2).

Three other mutations are placed on LGIII through their linkage to *Cbr-cnx-1*: *unc(sy5111)* is linked to it directly, and *dpy(sy5005)* and *sma(sa991)* through *unc(sy5111)* (Table 3). The *Cbr-dpy-1* allele *sy5022* maps to the left end of chromosome III by SNP-microarray mapping (Zhao *et al*. 2010).

*dpy(s1281)* was placed tentatively on LGIII by exclusion, because it showed no linkage to markers on LGI, II, IV, V and X (Table 2). One additional result supported this conclusion: a loose linkage to the LGIII sex-determination gene *Cbr-tra-1* (Kelleher *et al*. 2008). The molecular identification of *s1281* as an allele of *Cbr-eff-1*, which lies on chromosome II (3.50 Mb) (Lyu *et al*. 2026), does not support this assignment. We list the locus as *Cbr-eff-1* on LGII (Table 1). Although an earlier test with *Cbr-unc-4* (0.27 Mb) did not show linkage, a subsequent test with *Cbr-lin-31(sy5342)* (1.74 Mb) placed *Cbr-eff-1* 5.1 mu from *Cbr-lin-31* (Tables 2 and 3).

*dpy(sy5039)* and *dpy(sy5001)* are X-linked. They are linked to each other and to *unc(sa988)* in a three-factor cross (Table 4); *dpy(sy5001)* maps to the X chromosome by SNP-microarray and indel analysis (Koboldt *et al*. 2010; Zhao *et al*. 2010); and a *dpy(sy5039) dpy(sy5001)* double mutant was reported earlier as an X-chromosome marker (Inoue *et al*. 2007). *dpy(sy5057)* was also placed on X-chromosome based on a three-factor mapping involving a double mutant *dpy(sy5057) dpy(sy5001)* (Table 4). Eight other mutations, *unc(sy5077)*, *unc(sa988)*, *Cbr-unc-2(sy5329)*, *dpy(nm3)*, *dpy(sa971)*, *dpy(sa995)*, *lon(sy5227)* and *sma(sy5159)*, are placed on the X chromosome because F1 males from crosses of mutant hermaphrodites with wild-type males showed the mutant phenotype (Table 2).

**Table 4.**
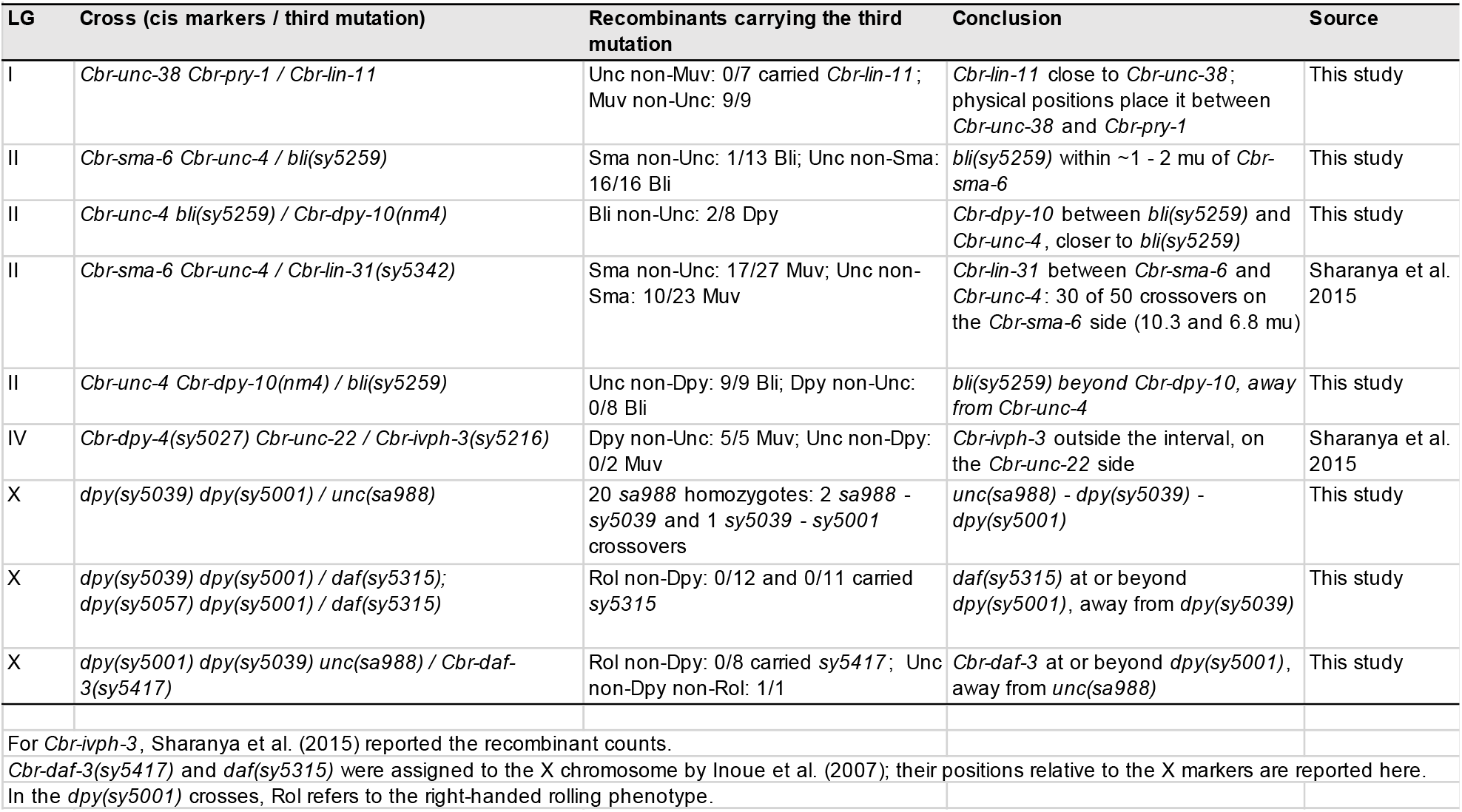
Three-factor crosses. Recombinant F2 animals of the stated phenotype were picked, and their progeny were scored for the third mutation.

### Two- and three-factor mapping and a genetic linkage map

Table 3 lists the two-factor crosses along with distances (Materials and Methods). Table 4 lists the three-factor crosses, and Figure 2 shows the linkage map.

**Figure 2.**
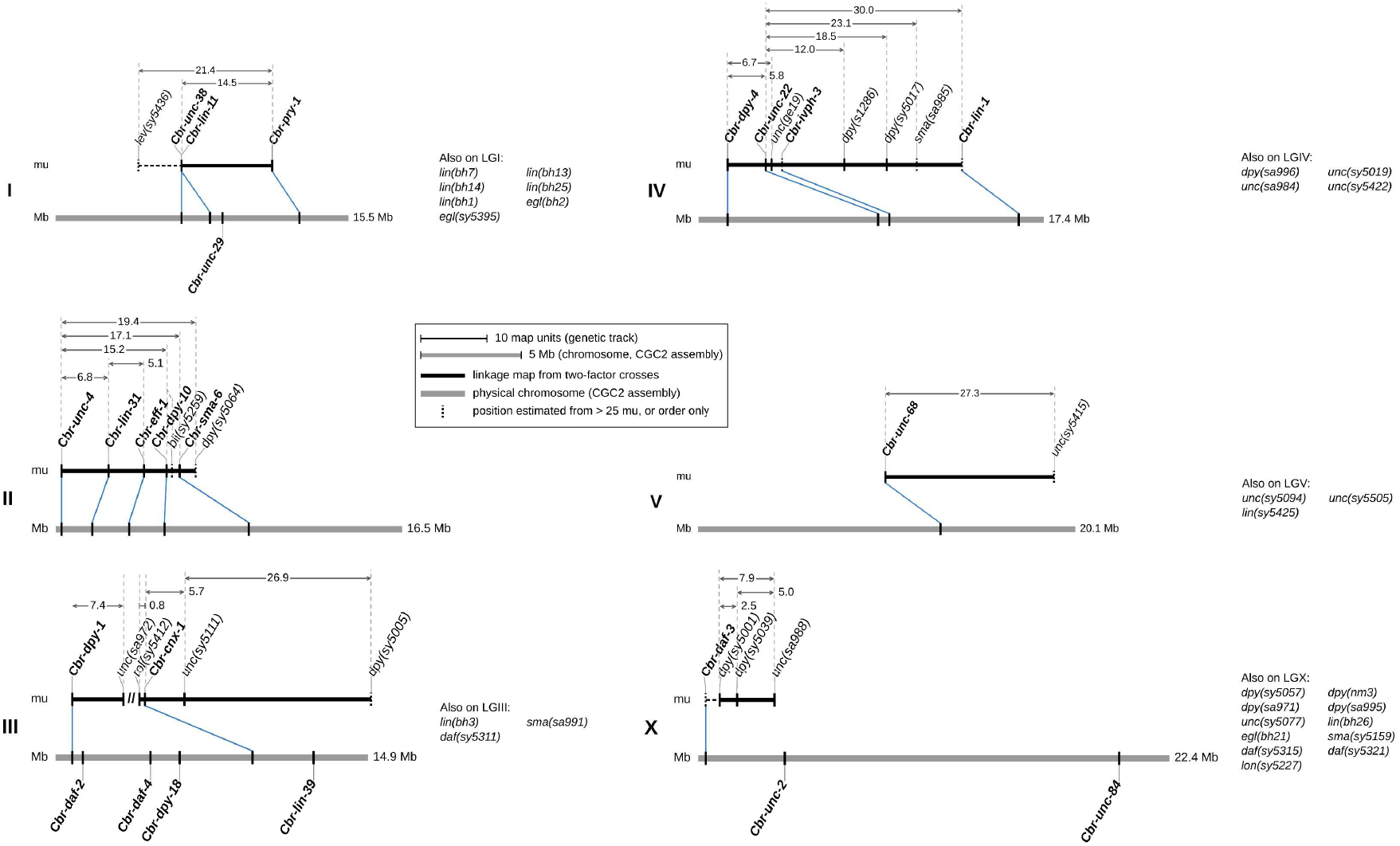
Genetic linkage map of *C. briggsae* based on mutants described in this study. The map is aligned with the physical chromosomes. For each linkage group, the upper (black) track is the genetic map from two-factor crosses (Table 3), drawn to scale in map units (mu), with the measured intervals shown by arrows above it. The lower (grey) bar is the corresponding chromosome of the CGC2 assembly, drawn to scale. Molecularly identified genes are in bold, and blue lines connect the genetic and physical positions of each gene. The genetic track covers only the interval spanned by crosses. Dotted ticks mark approximate positions: those estimated from clone tests or from long distances (> 25 mu) or three-factor crosses, those whose order relative to neighboring loci is not resolved, and those known only in order. A dashed line joins a locus whose side is not determined to the rest of the group. On LGI, *lev(sy5436)* lies 21.4 mu from *Cbr-pry-1*, but its side is not determined, and it is drawn on the *Cbr-unc-38* side. *Cbr-unc-29* is shown on the chromosome only, because its clone-test distance to *Cbr-pry-1* is not precise. On LGII, *Cbr-eff-1* is drawn on the *Cbr-sma-6* side of *Cbr-lin-31*, according to its physical position. The two LGIII blocks are not connected by any cross. The second block is anchored by *Cbr-cnx-1*, but its orientation is not known. On LGIV, *Cbr-ivph-3* is placed beyond *Cbr-unc-22* based on a three-factor cross without a distance (Table 4). The side of *Cbr-unc-22* on which *unc(ge19)*, *dpy(s1286)*, *dpy(sy5017)*, and *sma(sa985)* lie is not determined; they are drawn to the right. On LGV, *unc(sy5415)* lies 27.3 mu from *Cbr-unc-68* but its side is not determined, and it is drawn to the right. On the X chromosome, *Cbr-daf-3* orients the group. Genes with a physical position but no two-factor data are shown on the chromosome only. Loci and mutations assigned to a linkage group but not drawn on the map are listed on the right.

#### LGI

In a three-factor cross involving *Cbr-unc-38 Cbr-pry-1* double mutant and *Cbr-lin-11*, we scored Unc non-Muv and Muv non-Unc recombinants (Table 4), which suggested that *Cbr-lin-11* is close to *Cbr-unc-38*, on the side away from *Cbr-pry-1* or between the two. The physical positions of the three genes on chromosome I (6.65, 8.15 and 12.86 Mb) place *Cbr-lin-11* between the other two. In a four-class F2 cross, *Cbr-unc-38* and *Cbr-pry-1* are 14.5 mu apart (Table 3), and a clone test gave a consistent estimate (Supplementary File S1). No recombination was detected between *Cbr-unc-38* and *Cbr-lin-11* (0 of 42 clones; Table 3). *Cbr-unc-29(sy5437)* and *lev(sy5436)* lie 18.9 and 21.4 mu from *Cbr-pry-1*, respectively (Table 3).

#### LGII

Four mutations lie 15-19 mu from *Cbr-unc-4* (Table 3). Three-factor crosses place *bli(sy5259)* within 1-2 mu of *Cbr-sma-6*, and *Cbr-dpy-10(nm4)* between bli(sy5259) and *Cbr-unc-4*, close to *bli(sy5259)* (Table 4). *Cbr-dpy-10* and *Cbr-sma-6* are closely linked but, as expected for two different genes, complement each other (0/51 non-Unc cross progeny of *Cbr-sma-6 Cbr-unc-4* hermaphrodites and *nm4/+* males were Dpy). *Cbr-lin-31* lies between *Cbr-sma-6* and *Cbr-unc-4*, 10.3 mu from *Cbr-sma-6* and 6.8 mu from *Cbr-unc-4* (Sharanya *et al*. 2015) (Tables 3 and 4), in agreement with the physical positions of the three genes (*Cbr-unc-4* 0.27 Mb, *Cbr-lin-31* 1.74 Mb, and *Cbr-sma-6* 9.20 Mb). The physical position of *Cbr-eff-1* (3.50 Mb) places it on the *Cbr-sma-6* side. The five molecularly identified LGII genes, *Cbr-unc-4* (0.27 Mb), *Cbr-lin-31* (1.74 Mb), *Cbr-eff-1* (3.50 Mb), *Cbr-dpy-10* (5.18 Mb) and *Cbr-sma-6* (9.20 Mb), lie in an order consistent with the two- and three-factor crosses. Of the four mutations located 15-19 mu from *Cbr-unc-4* (Table 3), only the position of *dpy(sy5064)* could not be resolved because of differences between its two-factor and clone-test estimates (19.4 and 5.0 mu) (see Table 3 and File S1).

#### LGIII

Our mapping data for mutants support two separate linkage groups or blocks (Figure 2). In one of these, *Cbr-dpy-1* lies 7.4 mu from *unc(sa972)* (Table 3). Both map to the left arm of chromosome III (Koboldt *et al*. 2010), and *Cbr-dpy-1* lies at 0.79 Mb.

The other LGIII block spans at least 27 mu and contains *rol(sy5412)*, *Cbr-cnx-1*, *unc(sy5111)*, *dpy(sy5005)*, and *sma(sa991)* (Table 3). The order within the block is deduced from distances only, and *sma(sa991)* has no reliable distance. *Cbr-cnx-1* lies at 9.37 Mb on chromosome III (Lyu *et al*. 2026), which anchors this block far from *Cbr-dpy-1* (0.79 Mb). None of our existing genetic crosses connect the two blocks. *rol(sy5412)* was placed about 3 Mb from *Cbr-cnx-1* based on the SNP-microarray peak (at about 6.5 Mb, cb3) (Zhao *et al*. 2010), although the two are only 0.8 mu apart (Table 3). The difference probably reflects a lack of precision in the position of linkage peaks and the differences between the cb3 assembly and later assemblies (Ross *et al*. 2011; O’connor *et al*. 2026).

#### LGIV

Two-factor crosses link ten LGIV mutations into a single group (Table 3, Figure 2). *Cbr-unc-22*, *unc(ge19)*, *unc(sy5019)*, and *unc(sa984)* each lie within about 7 mu of *Cbr-dpy-4(sy5027)*. *dpy(s1286)* lies 12-14 mu from *Cbr-unc-22* and *unc(ge19)*, and *dpy(sy5017)* 18-26 mu from the cluster. *dpy(sa996)* is linked to the group, but its position is not determined (see above). The Muv gene *Cbr-lin-1(sa993)* lies 30.0 mu from *Cbr-unc-22*. *sma(sa985)* lies 23.1 mu from *Cbr-unc-22* (Table 3), probably on the *Cbr-lin-1* side. The physical positions of *Cbr-dpy-4* (1.47 Mb), *Cbr-unc-22* (9.05 Mb), *Cbr-*ivph*-3* (9.61 Mb), and *Cbr-lin-1* (16.14 Mb) orient the group and agree with the three-factor cross that places *Cbr-ivph-3* outside the interval between *Cbr-dpy-4* and *Cbr-unc-22*, on the *Cbr-unc-22* side (Sharanya *et al*. 2015) (Table 4, Figure 2). We have not determined which side of *Cbr-unc-22, dpy(s1286)* and *dpy(sy5017)* are located, and the order of *unc(ge19)*, *unc(sy5019),* and *unc(sa984)* relative to *Cbr-dpy-4* and *Cbr-unc-22*.

#### LGV

*Cbr-unc-68(sa997)* is the only molecularly identified gene on LGV (Lyu *et al*. 2026), which lies at 12.94 Mb on chromosome V. This physical location disagrees with earlier SNP-microarray mapping result that had placed the gene at about 7.2 Mb, and *unc(sy5094)*, *unc(sy5505),* and *unc(sy5415)* near 6.9, 8.5, and 15.1 Mb, respectively (Zhao *et al*. 2010; Sharanya *et al*. 2012). This means the cb3 peak for *sa997* is about 5.7 Mb from *Cbr-unc-68*. Linkage peaks can lie some distance from the gene (Zhao *et al*. 2010). Also, cb3 coordinates are not directly comparable with later assemblies, because the cb4 assembly moved, ordered, and reoriented large parts of the cb3 sequence (Ross *et al*. 2011). We therefore place *Cbr-unc-68(sa997)* on chromosome V based on its physical location and do not draw the cb3 peaks of any of the other three mutations (Figure 2).

*Cbr-unc-68(sa997)* is also linked to indels bhP24 and bhP31 (Koboldt *et al*. 2010), and *lin(sy5425)* to bhP5 (Sharanya *et al*. 2012). The linkage test of *sa997* and *unc(sy5415)* shows that the two mutations are linked and 27.3 mu apart (Tables 2 and 3), which confirms *sy5415* lies on the same chromosome as *Cbr-unc-68* and is consistent with the molecular position of the gene, but the side of *Cbr-unc-68* on which *unc(sy5415)* lies is not determined.

#### LGX

In a three-factor cross, *unc(sa988)* segregated from *dpy(sy5039)* and *dpy(sy5001)* in an order and frequency that place the three mutations as *sa988* - *sy5039* - *sy5001*. *dpy(sy5039)* lies 5.0 mu from *unc(sa988)* and 2.5 mu from *dpy(sy5001)*, and *unc(sa988)* and *dpy(sy5001)* are 7.9 mu apart (Tables 3 and 4).

The two *daf* genes, *daf(sy5315)* and *Cbr-daf-3(sy5417)*, that were assigned earlier to the X chromosome (Inoue *et al*. 2007), lie at or beyond *dpy(sy5001)*, on the side away from *unc(sa988)* (Table 4). Because *Cbr-daf-3* lies at 0.37 Mb, at the left end of the X chromosome, this orients the group, with *dpy(sy5001)* on the left (Figure 2). The SNP-microarray linkage peak of *dpy(sy5001)*, at about 7.2 Mb (cb3 assembly) (Zhao *et al*. 2010), lies to the right of *Cbr-daf-3*, in agreement with this orientation, although cb3 and CGC2 positions are not directly comparable (Ross *et al*. 2011). *unc(sy5077)* lies 13.7 mu from *dpy(sy5001)*, but the side of its location is not yet determined. *Cbr-unc-2(sy5329)* lies at 4.12 Mb (Lyu *et al*. 2026), however the gene has not been mapped relative to other X markers.

### Molecular identification of conserved genes

A total of 23 genes has been identified as orthologs of *C. elegans* genes (Table 5). Of these, 10 were characterized previously and include *Cbr-lin-11*, *Cbr-lin-39* and *Cbr-unc-84* (Sharanya *et al*. 2012); *Cbr-pry-1* (Seetharaman *et al*. 2010); *Cbr-lin-1*, *Cbr-lin-31* and *Cbr-ivph-3* (Sharanya *et al*. 2015; Chamberlin *et al*. 2020; Jhaveri *et al*. 2025); and *Cbr-daf-2*, *Cbr-daf-3* and *Cbr-daf-4* (Inoue *et al*. 2007). *Cbr-dpy-18(mf104)* is a *Mos1* insertion (see above) and another six were identified by sequencing in a parallel study (Lyu *et al*. 2026): *Cbr-dpy-4(sy5027)*, *Cbr-eff-1(s1281)*, *Cbr-dpy-10(nm4), Cbr-unc-68(sa997)*, *Cbr-cnx-1(sy5330),* and *Cbr-unc-2(sy5329)* (Table 5). For the remaining six, we report the supporting evidence here: *Cbr-dpy-1*, *Cbr-sma-6*, *Cbr-unc-4* and *Cbr-unc-22* below, and *Cbr-unc-38* and *Cbr-unc-29* with the levamisole-resistant mutants in the next section.

**Table 5.** Molecularly identified genes. Positions (Mb) are the lower gene coordinates in the CGC2 assembly for *C. briggsae*. The older cb4 assembly coordinates are also included for comparison.

| Gene | Alleles | LG | CGC2 Mb | cb4 Mb | Evidence | Source |
|---|---|---|---|---|---|---|
| *Cbr-unc-38* | *sy5440* | I | 6.65 | 6.69 | Fosmid rescue; missense mutation | This study |
| *Cbr-unc-29* | *sy5437* (+4) | I | 8.81 | 8.81 | Fosmid rescue; reduced transcript | This study |
| *Cbr-dpy-1* | *s1272*, *ge12*, *sy5016* (+20) | III | 0.79 | 0.78 | *dpy-1* cosmid M01E10 rescue | This study |
| *Cbr-sma-6* | *sy5148*, *sa983* | II | 9.2 | 9.42 | rescue of phenotype; missense mutation (sy5148) | This study; Lyu et al., 2026 |
| *Cbr-unc-4* | *sy5341* | II | 0.27 | 0.24 | Rescue by *C. elegans unc-4* | This study |
| *Cbr-unc-22* | *s1270* | IV | 9.05 | 9.00 | Twitching, nicotine response; fails to complement bh29; nonsense mutation | This study; Zhao et al. 2010; Lyu et al., 2026 |
| *Cbr-dpy-4* | *sy5027, sy5008* | IV | 1.47 | 1.43 | Nonsense mutation (*sy5027*) | Lyu et al., 2026 |
| *Cbr-eff-1* | *s1281* (+3) | II | 3.5 | 3.59 | Nonsense mutation (s1281) | Lyu et al., 2026 |
| *Cbr-dpy-10* | *nm4* | II | 5.18 | 5.28 | Nonsense mutation | Lyu et al., 2026 |
| *Cbr-cnx-1* | *sy5330, sy5002* | III | 9.37 | 9.26 | Missense mutation (sy5330) | Lyu et al., 2026 |
| *Cbr-unc-68* | *sa997* | V | 12.94 | 12.55 | Nonsense mutation | Lyu et al., 2026 |
| *Cbr-unc-2* | *sy5329* | X | 4.12 | 4.02 | Missense mutation | Lyu et al., 2026 |
| *Cbr-dpy-18* | *mf104* | III | 5.91 | 5.90 | Mos1 insertion | CGC (M.-A. Félix); Le et al. 2017 |
| *Cbr-lin-11* | *sy5336*, *sy5368* | I | 8.15 | 8.26 | Splice-site lesions; rescue | Sharanya et al. 2012 |
| *Cbr-pry-1* | *sy5270*, *sy5353*, *sy5411* | I | 12.86 | 12.64 | Splice-site (*sy5353*) and frameshift (*sy5411*) lesions; rescue | Seetharaman et al. 2010 |
| *Cbr-lin-31* | *sy5342* (+3) | II | 1.74 | 1.75 | Lesions (*gu138*, *gu162*) | Sharanya et al. 2015 |
| *Cbr-lin-39* | *bh20*, *bh23* | III | 12.29 | 11.93 | Point mutation and deletion; rescue | Sharanya et al. 2012 |
| *Cbr-lin-1* | *sa993*, *bh9*, *gu198* | IV | 16.14 | 16.41 | Nonsense mutation and deletion | Sharanya et al. 2015 |
| *Cbr*-ivph-3 | *sy5216*, *sy5392* | IV | 9.61 | 9.57 | Nonsense mutation (*sy5216*); rescue | Chamberlin et al. 2020; Jhaveri et al. 2025 |
| *Cbr-unc-84* | *sy5506* | X | 20.03 | 19.31 | Deletion; rescue | Sharanya et al. 2012 |
| *Cbr-daf-2* | *sy5445* | III | 1.3 | 1.28 | Missense, kinase domain | Inoue et al. 2007 |
| *Cbr-daf-3* | *sy5417* | X | 0.37 | 0.28 | Missense, MH2 domain | Inoue et al. 2007 |
| *Cbr-daf-4* | *sa973* | III | 4.52 | 4.45 | Missense, kinase domain; rescue of Sma | Inoue et al. 2007 |

#### Cbr-dpy-1

Several Dpy alleles, including *s1272, ge12*, and *sy5016*, fail to complement one another and define a locus that was first named *cby-4*. The gene maps to the left end of chromosome III (Koboldt *et al*. 2010; Zhao *et al*. 2010), where the *dpy-1* ortholog lies (III: 0.79 Mb). The locus has been called the *dpy-1* ortholog since 2007 (Inoue *et al*. 2007; Zhao *et al*. 2010; Sharanya *et al*. 2015), but no evidence was provided in those studies. The *C. elegans* cosmid M01E10, which carries *dpy-1* (M01E10.2), rescued the Dpy phenotype of *dpy(ge12)* in two independent transgenic lines. As expected, animals that lost the array were Dpy, confirming that the rescue depended on the transgene.

#### Cbr-sma-6

*sma(sy5148)* (originally *cby-15*) animals are small, and males have crumpled spicules and ray defects. A 6 kb *C. briggsae* genomic fragment containing *Cbr-sma-6* rescued both the body size and the male tail defects. *sy5148* maps to chromosome II by SNP-microarray and indel analysis (Koboldt *et al*. 2010; Zhao *et al*. 2010), and the SNP-microarray linkage peak lies about 0.6 Mb from *Cbr-sma-6* in the cb3 assembly (Zhao *et al*. 2010). *Cbr-daf-4* mutants are also small, but they form dauers constitutively (Inoue *et al*. 2007); *sy5148* animals do not form dauers, even at 27 °C or 30 °C (Inoue *et al*. 2007), as expected for a defect in *Cbr-sma-6* rather than *Cbr-daf-4*. A second allele, *sa983*, failed to complement *sy5148* (Supplementary File S1). Molecular analysis of *sy5148* has revealed a missense mutation in the kinase domain of the gene (Lyu *et al*. 2026).

#### Cbr-unc-4

*unc(sy5341)* animals cannot move backward, a phenotype that was rescued by a *C. elegans unc-4* genomic clone. The locus maps to LGII (Tables 2 and 3), consistent with the position of *Cbr-unc-4* at the left end of chromosome II (0.27 Mb), and it has served as the LGII reference marker in mapping studies (Koboldt *et al*. 2010; Sharanya *et al*. 2015). The precise mutation in *sy5341* has not been identified.

#### Cbr-unc-22

*s1270* and five other mutations, *sy5004*, *sy5053*, *sy5151*, *sy5168* and *sy5501*, cause twitching. Only *s1270* has been assigned to *Cbr-unc-22*, by complementation (below). The other five have not been tested against it, although their phenotype suggests that they are *Cbr-unc-22* alleles. Their twitching phenotypes are hallmarks of *unc-22* mutants in *C. elegans* (Moerman and Baillie 1979). The *Cbr-unc-22* locus was placed on chromosome IV by BAC rescue of an independently isolated allele, *st10417* (Zhao *et al*. 2010). Consistent with this identity, *s1270* failed to complement *Cbr-unc-22(bh29)*, a CRISPR allele generated in our lab (Culp *et al*. 2020) (14/35 Unc in F1 ♂ of *s1270* ⚥ crossed with *bh29/+* ♂). Molecular analysis of *s1270* has identified a nonsense mutation in the gene (Lyu *et al*. 2026).

### Levamisole-resistant mutants identify *Cbr-unc-38* and *Cbr-unc-29*

Levamisole is a nicotinic agonist that paralyzes worms by activating the levamisole-sensitive acetylcholine receptor of body-wall muscle (Lewis *et al*. 1980b; Richmond and Jorgensen 1999). In *C. elegans*, mutations affecting this receptor cause an Unc phenotype and confer strong resistance to the chemical (Lewis *et al*. 1980a; Fleming *et al*. 1997). The *C. briggsae lev* mutations were isolated in two different screens (Materials and Methods). All Lev mutant strains are Unc, and none of the five tested is X-linked (Table 2). Complementation tests define at least three distinct loci (Figure 3A, Table 6). Five alleles, *sy5437*, *sy5439*, *sy5502*, *sy5503* and *sy5504*, fail to complement one another in every combination tested and form a single group. *sy5440* and *sy5436* complement this group and each other, and each defines a separate locus. *sy5438* complements the *sy5437* group and *sy5436* but has not been tested against *sy5440*. It may therefore be an allele of *sy5440* or define a fourth locus.

**Figure 3.**
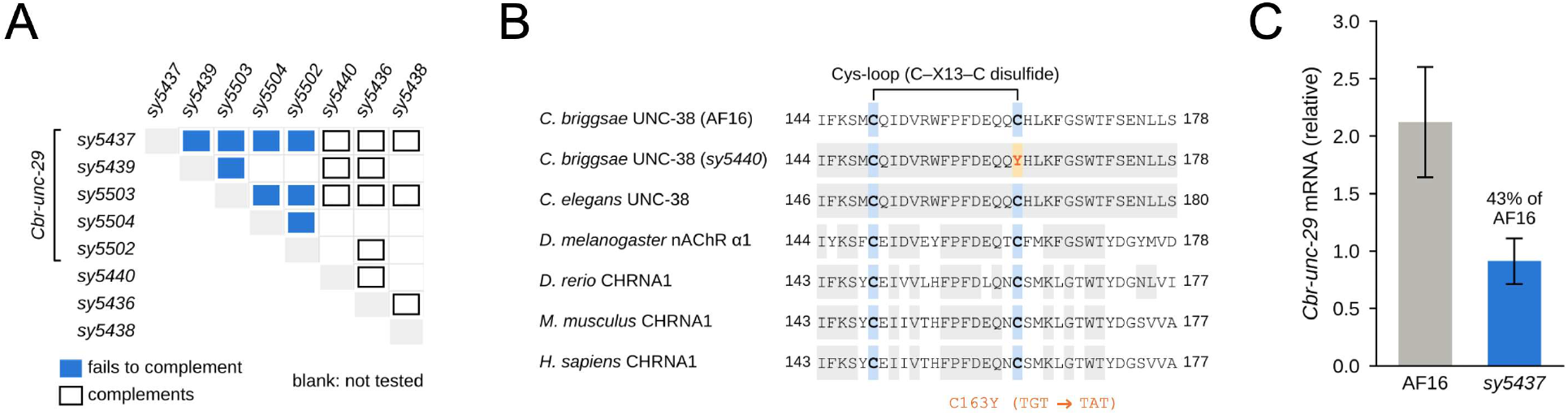
Levamisole-resistance genes. (A) Complementation tests among eight *lev* mutant alleles. Filled squares, failure to complement; open squares, complementation; blank, not tested. The five *Cbr-unc-29* alleles are indicated. (B) Alignment of the Cys-loop region of UNC-38 from *C. briggsae* AF16, *lev(sy5440)*, and *C. elegans* with the nicotinic acetylcholine receptor α1 subunits of *D. melanogaster*, zebrafish, mouse, and human. The two cysteines that form the Cys-loop disulfide are shaded blue, residues identical to *C. briggsae* UNC-38 are shaded grey, and the *sy5440* substitution (C163Y; TGT to TAT) is highlighted. (C) qRT-PCR analysis of *Cbr-unc-29* transcripts in AF16 and *lev(sy5437)*. Mean ± SD of three biological replicates is plotted.

**Table 6.**
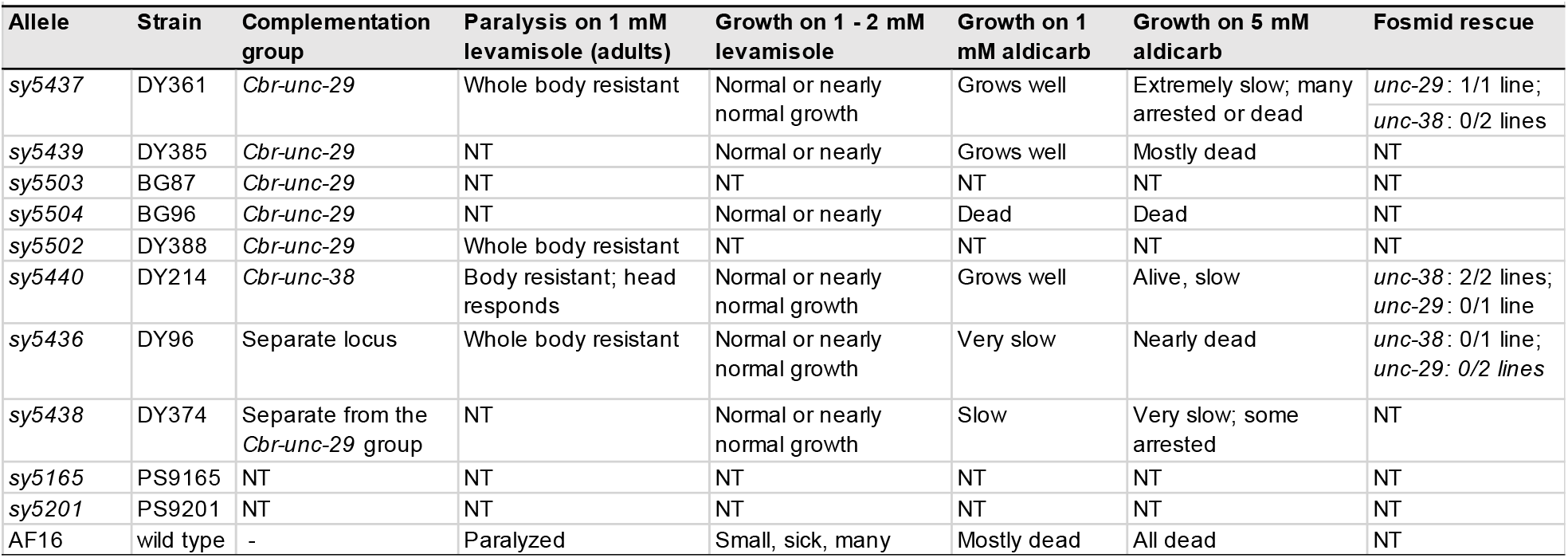
Levamisole-resistant mutants. Complementation, responses to drugs, and rescue are shown. Paralysis was scored in adults placed on 1 mM levamisole; growth was scored qualitatively after several days on drug plates (Materials and Methods). NT, not tested.

| Allele | Strain | Complementation group | Paralysis on 1 mM levamisole (adults) | Growth on 1 - 2 mM levamisole | Growth on 1 mM aldicarb | Growth on 5 mM aldicarb | Fosmid rescue |
|---|---|---|---|---|---|---|---|
| *sy5437* | DY361 | *Cbr-unc-29* | Whole body resistant | Normal or nearly normal growth | Grows well | Extremely slow; many arrested or dead | *unc-29*: 1/1 line;<br>*unc-38*: 0/2 lines |
| *sy5439* | DY385 | *Cbr-unc-29* | NT | Normal or nearly | Grows well | Mostly dead | NT |
| *sy5503* | BG87 | *Cbr-unc-29* | NT | NT | NT | NT | NT |
| *sy5504* | BG96 | *Cbr-unc-29* | NT | Normal or nearly | Dead | Dead | NT |
| *sy5502* | DY388 | *Cbr-unc-29* | Whole body resistant | NT | NT | NT | NT |
| *sy5440* | DY214 | *Cbr-unc-38* | Body resistant; head responds | Normal or nearly normal growth | Grows well | Alive, slow | *unc-38*: 2/2 lines;<br>*unc-29*: 0/1 line |
| *sy5436* | DY96 | Separate locus | Whole body resistant | Normal or nearly normal growth | Very slow | Nearly dead | *unc-38*: 0/1 line;<br>*unc-29: 0/2 lines* |
| *sy5438* | DY374 | Separate from the *Cbr-unc-29* group | NT | Normal or nearly normal growth | Slow | Very slow; some arrested | NT |
| *sy5165* | PS9165 | NT | NT | NT | NT | NT | NT |
| *sy5201* | PS9201 | NT | NT | NT | NT | NT | NT |
| AF16 | wild type | - | Paralyzed | Small, sick, many | Mostly dead | All dead | NT |

*lev(sy5440)* maps to chromosome I by SNP-microarray mapping (Zhao *et al*. 2010). It is linked to *Cbr-pry-1* and *Cbr-lin-11* (Table 2) but not to markers on LGII, LGIII, LGIV or the X chromosome (Table 2). *lev(sy5437)* is also linked to *Cbr-pry-1*, and *lev(sy5439)* is unlinked to markers on LGII and LGIV (Table 2). *lev(sy5436)* is linked to *Cbr-pry-1* (21.4 mu; Table 3) but not to *Cbr-sma-*6 or *Cbr-dpy-4* (Table 2), which places it on LGI.

The mutants also differ in their responses to drugs (Table 6). On plates containing 1 or 2 mM levamisole, where AF16 animals were small, sick and often died, the six mutants tested grew normally or nearly normally. Aldicarb, an acetylcholinesterase inhibitor, kills wild-type animals through excess acetylcholine, and mutants with reduced acetylcholine release show resistance (Miller *et al*. 1996). On 1 mM aldicarb containing plates most AF16 animals died and *lev* mutants showed different responses. Although *sy5437*, *sy5439* and *sy5440* grew normally, *sy5436* and *sy5438* grew slowly, and *sy5504* died. On 5 mM aldicarb, *sy5440* animals were alive but slow, whereas animals of the other strains were arrested, mostly dead or dead. Aldicarb resistance is therefore partial and varies among strains, including among the alleles of the *sy5437* group. Allelic series are common among *C. elegans lev* mutants (Fleming *et al*. 1997), but to our knowledge allele-specific differences in aldicarb response have not been reported. When exposed to 1 mM levamisole, *sy5436*, *sy5437*, and *sy5502* showed resistance in the whole body, but the heads of *sy5440* animals still responded. The head response therefore distinguishes *sy5440* from *sy5436* and the *sy5437* group. In *C. elegans*, *unc-38* mutants show the same pattern, with body muscles being more resistant to cholinergic agonists than head muscles (Fleming *et al*. 1997).

Because the *C. elegans unc-29* and *unc-38* genes encode non-α and α subunits of the receptor and lie on chromosome I (Fleming *et al*. 1997), we tested *C. elegans* fosmids carrying each gene for rescue of *sy5440*, *sy5437* and *sy5436* (Tables 5 and 6). The *unc-38* fosmid restored levamisole sensitivity to *lev(sy5440)* but not to *lev(sy5437)* or *lev(sy5436)*. The *unc-29* fosmid restored sensitivity to *lev(sy5437)* but not to *lev(sy5440)*. The *lev(sy5436)* lines carrying the *unc-29* fosmid were not rescued.

Sequencing of all eight *Cbr-unc-38* exons in *lev(sy5440)* identified a single G-to-A transition in exon 4, read on both strands, which converts codon 163 from TGT (cysteine) to TAT (tyrosine) (Figure 3B). Cys163 is the second cysteine of the Cys-loop, the 13-residue disulfide-bonded loop that defines the Cys-loop family of ligand-gated ion channels. Both cysteines and the FPFD motif within the loop are identical in *C. briggsae* and *C. elegans* UNC-38. Both cysteines and the FPFD motif within the loop are also conserved in the *Drosophila* nicotinic receptor α1 subunit and in the muscle receptor α1 subunits of zebrafish, mouse, and human (Figure 3B). In the vertebrate subunits, the residue equivalent to Cys163 is Cys142 of the mature protein (Cys162 of the precursor). A tyrosine at this position cannot form the disulfide bond. The coding exons of *Cbr-unc-29* sequenced in *lev(sy5440)* were wild-type. Together, the rescue, complementation and sequence data identify *lev(sy5440)* as an allele of *Cbr-unc-38*.

The *lev(sy5437)* animals showed no mutations in any of the 11 *Cbr-unc-29* exons. However, *Cbr-unc-29* transcript levels in *sy5437* animals are reduced (43% of AF16 controls; 0.91 ± 0.20 vs 2.12 ± 0.48; Figure 3C). Rescue by the *unc-29* fosmid, together with reduced transcript levels, suggests that *sy5437* is an allele of *Cbr-unc-29*, possibly affecting one of the introns or flanking sequences. More experiments are needed to identify the exact site of mutation in the gene.

### Comparison with *C. elegans*

All 23 molecularly identified *C. briggsae* genes lie on the same chromosome as their *C. elegans* orthologs (Table 5, Figure 4). This agrees with the conservation of chromosome identity between the two species (Stevens *et al*. 2022; O’connor *et al*. 2026). Gene order within chromosomes, however, differs. *Cbr-unc-4* and *Cbr-sma-6* are 17.1 mu apart in *C. briggsae* and lie at opposite ends of a 9 Mb interval on chromosome II, but their *C. elegans* orthologs are only about 2 mu apart. *Cbr-lin-39* lies on the right arm of chromosome III, whereas *C. elegans lin-39* is near the center of chromosome III, and *Cbr-lin-11* lies between *Cbr-unc-38* and *Cbr-unc-29*, whereas *C. elegans lin-11* lies beyond *unc-29*. On the X chromosome, by contrast, *Cbr-daf-3*, *Cbr-unc-2,* and *Cbr-unc-84* lie in the same order as their *C. elegans* orthologs. On chromosome IV, the four identified genes lie in exactly the reverse order of their *C. elegans* orthologs (Figure 4).

**Figure 4.**
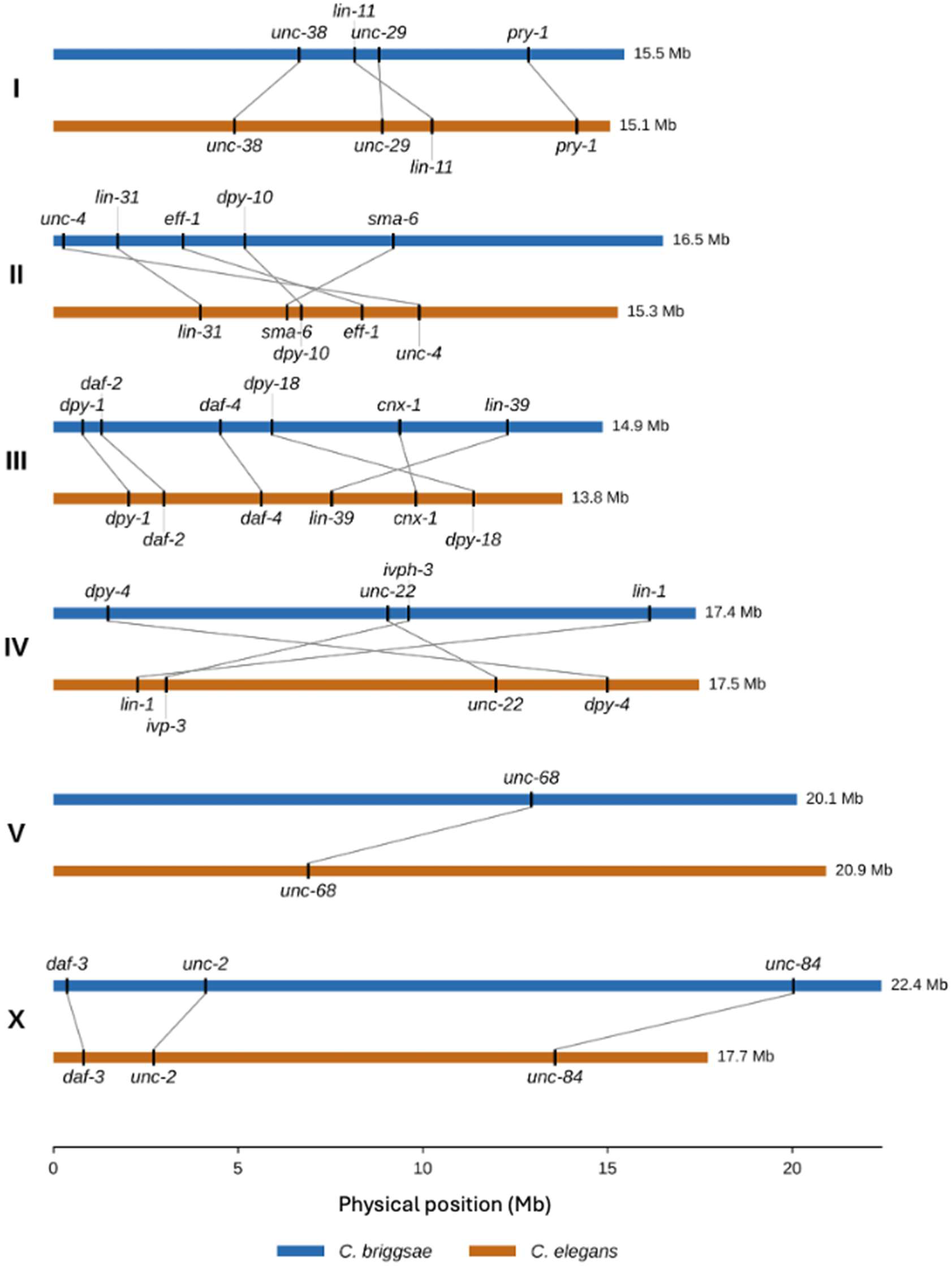
Positions of molecularly identified genes in *C. briggsae* (CGC2 genome assembly) and *C. elegans* (WBcel235 genome assembly). Bars are drawn to the length of each chromosome, and lines connect orthologs. Chromosomes are drawn in the orientation of each assembly. All genes lie on the same chromosome in the two species, but their order within chromosomes differs. On chromosome IV, the order of all four genes is reversed.

The Lev class offers a direct comparison of forward genetic analysis. In the *C. elegans* screen, more than 200 strongly resistant mutations fell into six genes: *unc-29*, *unc-63*, *unc-38*, *unc-74*, *unc-50*, and *lev-1* (Lewis *et al*. 1980a). Four of these genes (*unc-29*, *unc-38*, *unc-63,* and *unc-74*) lie on chromosome I. The two *C. briggsae* receptor genes identified here, *Cbr-unc-38* and *Cbr-unc-29*, are also on LGI, as is the third Lev locus, *lev(sy5436)*. As in *C. elegans*, *Cbr-unc-29* is the Lev gene with the most alleles in our collection.

Some phenotypic similarities are worth mentioning. *unc(sy5422)* animals are small, nearly paralyzed and dauer-defective, like *C. elegans unc-119* mutants. However, *sy5422* animals carry no mutation in the *Cbr-unc-119* coding region or its flanking sequences, and they are not rescued by *C. elegans* or *C. briggsae unc-119* clones (Z. Zhao, unpublished data). *sy5422* also maps to chromosome IV, whereas *C. elegans unc-119* is on chromosome III. *sy5422* animals did not survive ivermectin treatment, whereas *C. elegans unc-119* animals can tolerate it. Since ivermectin resistance in *C. elegans* is a feature of mutants with defective sensory cilia (Dent *et al*. 2000), this suggests that *sy5422* does not disrupt amphid cilia.

## Discussion

*C. briggsae* is a leading nematode system for developmental, physiological, and behavior studies, yet its forward genetics has relied on a handful of previously reported markers with easily observed phenotypes whose supporting data were not published in detail. This study provides that foundation. We describe 183 mutations, 121 of which define 55 loci, with the complementation tests, linkage tests and two-and three-factor crosses that place 51 loci and 15 further mutations on the six linkage groups. Further, we report genetic, rescue, and sequence evidence for the identities of six genes, including four (*Cbr-dpy-1*, *Cbr-sma-6*, *Cbr-unc-4*, and *Cbr-unc-22*) that have long served as mapping markers. Loci with two-factor positions (Table 3, Figure 2) are the most useful markers, because their distances to neighboring loci are known. Our work has identified such markers on all six linkage groups, for example *Cbr-unc-4*, *Cbr-sma-6*, and *Cbr-dpy-10* on LGII and *Cbr-unc-22*, *Cbr-dpy-4*, and *dpy(s1286)* on LGIV. On LGI, *Cbr-unc-38* and *Cbr-pry-1* are 14.5 mu apart, with *Cbr-lin-11* close to *Cbr-unc-38*. Earlier, Wei *et al*. (2014) noted that LGI, LGV and the X chromosome lacked sets of mapped markers and generated TALEN alleles of conserved genes. Our collection complements those alleles, notably on the X chromosome, where *unc(sa988)*, *dpy(sy5039)*, and *dpy(sy5001)* form a mapped cluster oriented by *Cbr-daf-3*. LGV remains poorly marked. While *Cbr-unc-68* anchors it physically, its only two-factor distance, 27.3 mu to *unc(sy5415)*, is long and rests on a small clone test. Together with data in Supplementary File S1, these strains can facilitate genetic mapping, strain construction, and comparative genetics in *C. briggsae*.

All 23 molecularly identified genes lie on the same chromosome in *C. briggsae* and *C. elegans*, consistent with genome-wide comparisons (Stevens *et al*. 2022; O’connor *et al*. 2026), but gene order within chromosomes differs (Figure 4). For positional cloning, synteny therefore predicts the chromosome on which a *C. briggsae* ortholog lies but not its position within that chromosome, and candidate genes should be ranked by their physical position in the *C. briggsae* assembly.

Our study presents first detailed analysis of levamisole-resistant mutants in *C. briggsae*. We found that *Cbr-unc-38(sy5440)* mutation removes a cysteine that is invariant across the Cys-loop receptor family and is expected to disrupt folding or assembly of the receptor. The parallel with *C. elegans*, where four of six Lev genes lie on chromosome I, extends to the third locus *lev(sy5436)*, which also maps to LGI, complements both receptor groups, and is not rescued by either fosmid, thereby making *Cbr-unc-63* and *Cbr-unc-74*, the other two *C. elegans* Lev genes on chromosome I, promising candidates.

More broadly, most loci reported here remain unidentified, four loci and 47 mutations are not yet assigned to a linkage group, and 62 mutations await complementation tests that would place them in loci. For some of these, map position and phenotype point to candidate genes. The five unmapped twitchers, *unc(sy5004)*, *unc(sy5053)*, *unc(sy5151)*, *unc(sy5168)*, and *unc(sy5501)*, are likely *Cbr-unc-22* alleles (Supplementary File S1), which can be confirmed by complementation tests with *Cbr-unc-22(s1270)*. The Rol mutation, *rol(sy5412)*, lies 0.8 mu from *Cbr-cnx-1* (9.37 Mb on chromosome III). No ortholog of a classical *C. elegans* Rol or Sqt gene and no named cuticle collagen gene is annotated in the nearby region of chromosome III. The nearest genes are *Cbr-col-105* (8.33 Mb) and *Cbr-col-91* (8.17 Mb). Because many *C. briggsae* collagen genes are annotated without names, whole-genome sequencing is likely to be the more efficient way to determine the molecular identity of *rol(sy5412)*. By contrast, *bli(sy5259)* lies 1-2 mu from *Cbr-sma-6* (9.20 Mb on chromosome II) on the *Cbr-unc-4* side (Table 4), between *Cbr-dpy-10* (5.18 Mb) and *Cbr-sma-6*, consistent with the position of *Cbr-bli-1* (7.53 Mb) but not *Cbr-bli-2* (9.85 Mb), which lies on the other side of *Cbr-sma-6*. In *C. elegans*, loss of either of these strut collagen genes causes blisters (Page and Johnstone 2007; Adams *et al*. 2023). Sequencing of the *Cbr-bli-2* coding region in *bli(sy5259)* animals revealed no mutation (data not shown), making *Cbr-bli-1* a strong candidate. Similarly, the small, nearly paralyzed and dauer-defective

*unc(sy5422)* animals resemble *C. elegans unc-33* and *unc-44* mutants, which are defective in axon outgrowth and guidance (Hedgecock *et al*. 1985; Li *et al*. 1992; Otsuka *et al*. 1995). *Cbr-unc-33* and *Cbr-unc-44* both lie on chromosome IV, and *Cbr-unc-44* (6.59 Mb) is close to the region to which *unc(sy5422)* maps, so sequencing or transgenic rescue could determine whether *sy5422* affects either ortholog. Dye filling would provide an independent test, since in *unc-33* and *unc-44* mutants the amphid and phasmid neurons fill normally with lipophilic dyes but project misrouted axons (Hedgecock *et al*. 1985). For mapped loci without an obvious candidate, whole-genome sequencing of mutants with known map intervals (e.g., see (Chamberlin *et al*. 2020)) can help determine their molecular identities. We hope that the strains and data presented here will make *C. briggsae* genetics more accessible to the community.

## ACKNOWLEDGMENTS

Genetic screens, mutant isolation, and mapping were assisted by many members of the Sternberg, Baillie, and Gupta laboratories, including Keith Brown, Michael Cho-Young, Iqra Effendi, Zobia Jawed, Prateek Goyal, Greta Jo, Jennifer Li, Sujatha Marri, Shirley Poon, Sophie Rangarajan, and Mohamad Zubair. We are grateful to many colleagues including Eric Haag, Marie-Anne Félix, Helen Chamberlin, and James Thomas for mutant alleles and some of data and strains reported here; David Miller for the *unc-4* plasmid pNC4-21; Raymond Miller for SNP genotyping; and Zhongying Zhao for the analysis of *Cbr-unc-119* and for SNP-microarray mapping. This work was supported by a Natural Sciences and Engineering Research Council of Canada (NSERC) Discovery grant and McMaster University funds to BPG and by the Howard Hughes Medical Institute, with which PWS was an Investigator. The authors declare no conflicts of interest.

## Supplementary Materials

**Supplementary File S1.** File containing all loci (with phenotypes and the evidence that defines each locus), alleles and strains, former names, linkage tests, mapping, transgenic lines, and molecular analyses (Microsoft Excel).

**Supplementary File S2.** Python script for calculating map distances and 95% confidence intervals.

**Table S1.** Primers used for sequencing and qRT-PCR (Microsoft Excel).

**Table S2.** Publication status of each locus and unassigned mutations (Microsoft Excel).

